# Unexpected redundancy of Gpr56 and Gpr97 during hematopoietic cell development and differentiation

**DOI:** 10.1101/2020.09.18.303230

**Authors:** A. Maglitto, S.A. Mariani, E. de Pater, C. Rodriguez-Seoane, C.S. Vink, X. Piao, M.-L. Lukke, E. Dzierzak

## Abstract

Integrated molecular signals regulate cell fate during embryonic hematopoietic stem cell (HSC) generation. The G-protein coupled receptor 56 (Gpr56) is the most highly-upregulated receptor gene in cells that take on hematopoietic fate and it is expressed by adult bone marrow HSCs. Although Gpr56 is required for hematopoietic stem/progenitor cell (HS/PC) generation in zebrafish embryos, its function in mammalian hematopoiesis remains unclear. Here we examine the role of Gpr56 in HS/PC development in Gpr56 conditional knockout (cKO) mouse embryos and Gpr knockout (KO) embryonic stem cell (ESC) hematopoietic differentiation cultures. Our results show a myeloid bias of Gpr56 cKO fetal liver HSCs and an increased definitive myeloid progenitor cell frequency in Gpr56KO ESC differentiation cultures. Surprisingly, we find that mouse Gpr97 rescues Gpr56 morphant zebrafish hematopoietic generation, and that Gpr97 expression is upregulated in mouse Gpr56 deletion models. When both Gpr56 and Gpr97 are deleted in ESCs, no/few HS/PCs are generated upon ESC differentiation. Together, our results reveal novel and redundant functions for these two G-protein coupled receptors in normal mammalian hematopoietic cell development and differentiation.

## Introduction

Hematopoietic stem cells (HSCs) are rare, self-renewing cells that sustain lifelong blood production and used for therapeutic reconstitution of the blood system. To improve such therapies, research has focused on identifying the signalling molecules directing the development of the first adult-repopulating HSCs. In vertebrate embryos, HSCs arise in the aorta-gonad mesonephros (AGM) region from hemogenic endothelial cells (HECs). HECs undergo a highly-conserved ‘endothelial-to-hematopoietic transition’ (EHT) process(Ciau-Uitz & Patient, 2019, Dzierzak & Bigas, 2018, Dzierzak & Speck, 2008, Jaffredo, Nottingham et al., 2005). Transcriptomics and loss-of-function studies have identified candidate regulators/signaling pathways in transitioning cells(Gao, Chen et al., 2020, Li, Gao et al., 2017, Lichtinger, Ingram et al., 2012, Moignard, Woodhouse et al., 2015, Porcheri, Golan et al., 2020, Swiers, Baumann et al., 2013, Zhou, Li et al., 2016).

We identified *Gpr56* as the highest-upregulated receptor gene during mouse EHT and localized its expression to emerging hematopoietic cells(Solaimani Kartalaei, Yamada-Inagawa et al., 2015). Gpr56 belongs to the adhesion G-protein coupled receptor family which mediates cell-cell and cell-matrix interactions(Langenhan, Aust et al., 2013). It is well-conserved across vertebrate species and contains four domains (external-N-terminal, GPCR-Autoproteolysis-INducing, 7-transmembrane, C-terminal-cytoplasmic)(Zhu, Luo et al., 2019). Upon ligand-binding, an auto-proteolytic cleavage divides the protein in two non-covalently linked fragments. The cytoplasmic domain signals through RhoA/ ROCK to trigger downstream cellular functions(Ackerman, Garcia et al., 2015, Luo, Jeong et al., 2014, Olaniru, Pingitore et al., 2018).

Gpr56 functions in several developmental processes and its dysfunction is linked to human genetic disorders. *Gpr56* deletion in the mouse causes a severe reduction in CNS myelination(Ackerman et al., 2015, Ackerman, Luo et al., 2018) and mutations in human GPR56 cause a recessive brain malformation, defective cerebral cortex and CNS hypomyelination(Cauley, Hamed et al., 2019, Piao, Hill et al., 2004). Interestingly, aberrant expression of GPR56 in leukemic stem cells is associated with high-risk/treatment-resistance in AML patients(Barjesteh van Waalwijk van Doorn-Khosrovani, Erpelinck et al., 2003, Daria, Kirsten et al., 2016, Groschel, Sanders et al., 2014, Pabst, Bergeron et al., 2016). Although Gpr56 is well-characterized in CNS development, its role in normal hematopoiesis is uncertain.

*Gpr56* knockdown zebrafish embryos suffer dramatic reductions in aortic HS/PC (hematopoietic stem/progenitor cell) generation during EHT(Solaimani Kartalaei et al., 2015). This deficiency could be rescued by zebrafish *gpr56* (also mouse *Gpr56*) mRNA injection. Thus, *gpr56* is necessary for zebrafish aortic HS/PC development(Solaimani Kartalaei et al., 2015). In contrast, although *Gpr56* is highly-expressed by mouse HS/PCs(Rao, Marks-Bluth et al., 2015), one mouse germline *Gpr56* (exon2-3) deletion study showed surprisingly small changes in HPC numbers and HSC-repopulating activity(Rao et al., 2015), whereas another(Saito, Kaneda et al., 2013) showed reduced HSC self-renewal. As some Gpr56 protein was detected in this mouse model (possibly a failure to delete the S4 splice variant(Rao et al., 2015)), a role for Gpr56 in normal mammalian hematopoietic development awaits further study.

Here we examine the role of Gpr56 *in vivo* and *in vitro* during mouse hematopoietic development. We show that *in vivo* transplanted *Gpr56* conditional knockout mouse fetal liver (FL) HSCs and *Gpr56*^*-/-*^ mouse embryonic stem cell (ESC)-derived HPCs are myeloid-biased, that mouse *Gpr97* mRNA can rescue HS/PC generation in *gpr56* morphant zebrafish embryos, and that deletion of *Gpr56* in mouse FL HSCs and ESC-derived HPCs results in upregulated *Gpr97* expression. Deletion of both *Gpr56* and *Gpr97* in mouse ESC results in almost complete loss of HPC production, thus revealing previously unrecognized and important redundant roles for these two GPCRs in mouse hematopoietic development and differentiation.

## Results

### Yolk sac hematopoietic progenitors are reduced in *Gpr56* conditionally-deleted embryos

A conditional knockout (cKO) approach was taken to delete all isoforms of *Gpr56* in mouse embryonic cells expressing *VEC(vascular endothelial-cadherin)* or *Vav* prior to and following HSC generation, respectively. Yolk sacs (YS) isolated from embryonic day (E)9*VECCre:loxGpr56* cKO and control wild type (WT) littermates (**Fig1A**) contained equivalent cell numbers **(Fig EV1A**). CD31^+^ cKO YS cells showed a significant decrease in relative *Gpr56* mRNA as compared to WT controls (**Fig1B**) and *Gpr56* deletion was verified by DNA-PCR (**Fig EV1B, upper**). *In vitro* colony forming unit-culture (CFU-C) assays revealed a significantly lower number of HPCs (CFU-C/YS) in the cKO as compared to WT (**Fig1C**). *Gpr56* recombination was verified in 18/19 individual cKO colonies (**Fig EV1B, lower**). Flow cytometry revealed a significantly decreased cKit^+^ percentages in cKO YS as compared to WT, but only slight reductions in erythro-myeloid (EMPs; CD41^+^cKit^+^CD16/32^+^) and other (CD31^+^cKit^+^, CD45^+^cKit^+^) progenitor percentages (**Fig EV1C, Fig1D**).

**Figure 1.**
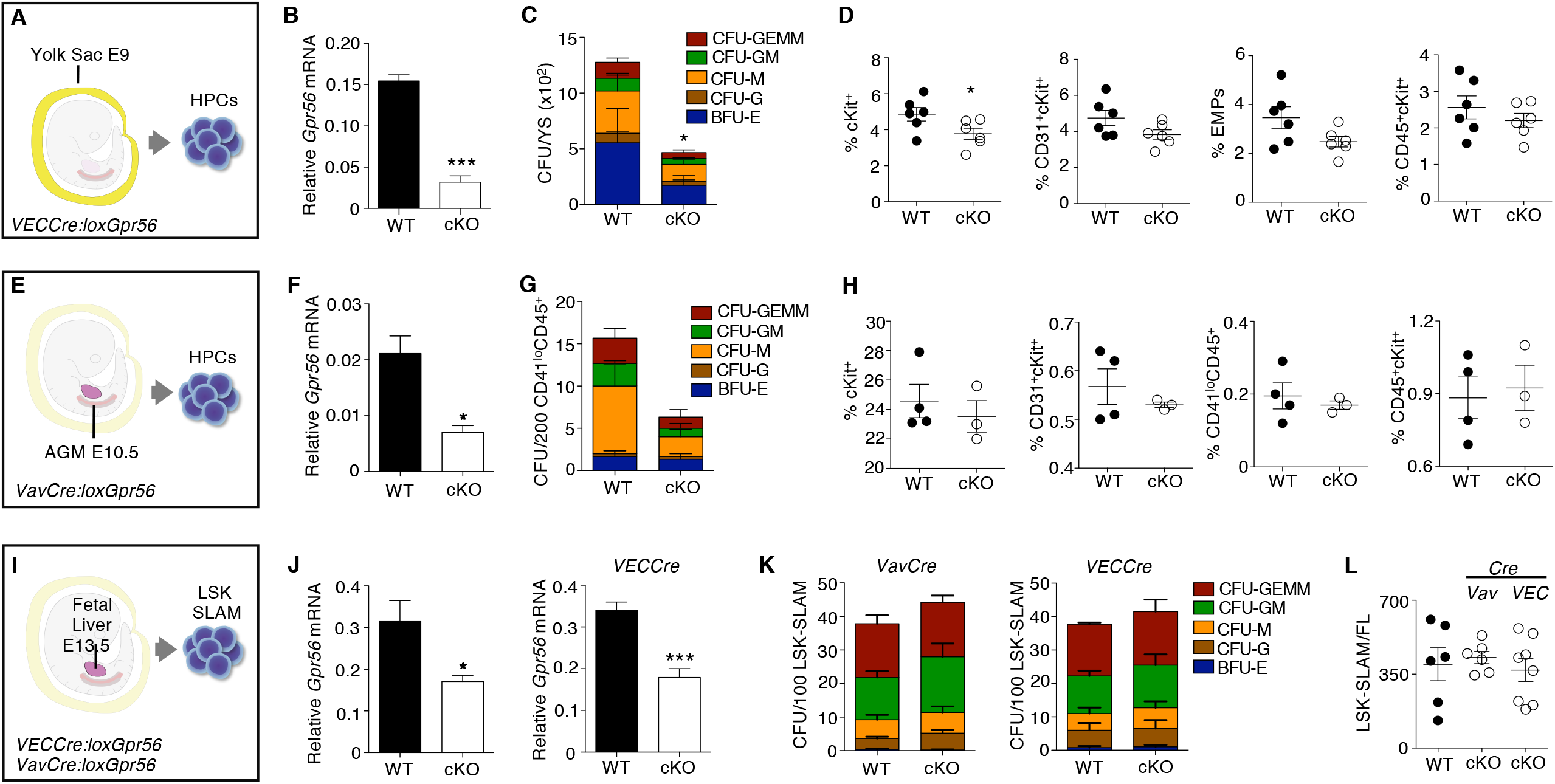
Gpr56 deficiency effects early hematopoietic development in mouse embryos. Experimental setup for **A)** E9 yolk sac (YS, 21-25 somite pairs), **E)** E10.5 AGM (34-37sp) and **I)** E13.5 fetal liver (FL) hematopoietic progenitor cell (HPC) analyses. **B)** Relative expression of *Gpr56* in wild type (WT) and *VECCre:loxGpr56* conditional knockout (cKO) E9 YS cells normalized to *β-actin* by RT-PCR analysis (n=3). **C)** Number of colony forming units (CFU) per WT and cKO E9 YS (n=3). **D)** Percentages of cKit^+^, CD31^+^cKit^+^,CD41^+^cKit^+^CD16/32^+^ (EMP=erythro-myeloid progenitor), and CD45^+^ckit^+^ cells in WT and cKO E9 yolk sacs (n=6). Flow cytometric gating strategy shown in Fig EV2. **F)** Relative expression of *Gpr56* in wild type (WT) and *VavCre:loxGpr56* conditional knockout (cKO) E10.5 AGM cells normalized to *β-actin* by RT-PCR analysis (n=3). **G)** Number of colony forming units (CFU) per WT and cKO E10.5 AGM (n=3). **H)** Percentages of cKit^+^, CD31^+^cKit^+^, CD41^lo^CD45^+^ and CD45^+^ckit^+^ cells in WT and cKO E10.5 AGMs (n=6). **J)** Relative expression of *Gpr56* in WT and *VavCre:* and *VECCre:loxGpr56* cKO LSK-SLAM sorted E13.5 FL cells normalized to *β-actin* by RT-PCR analysis (n=5). *p≤0.05. **K)** Number of CFU per WT and *Vav* and *VECCre* cKO E13.5 FL LSK-SLAM cells (n=4). **L)** Number of LSK-SLAM cells per WT and *Vav* and *VECCre* cKO E13.5 FL (n=6). Distinct colony types are indicated. Distinct colony types are indicated. CFU-GEMM=granulocyte, erythroid, macrophage, megakaryocyte; -GM=granulocyte, macrophage; -M=macrophage; G=granulocyte; BFU-E=burst forming unit-erythroid. Mean±SEM are shown.

Phenotypic analysis of AGM cells from the same cohort of E9 *VECCre:loxGpr56* cKO embryos showed a significant decrease in the percentage of cKit^+^ cells and slight reductions in EMPs and other progenitors (**Fig EV1D)**. In contrast, no reductions in percentages of cKit^+^, EMPs or other progenitor cells were found in E10.5 *VECCre:loxGpr56* cKO AGM cells (**Fig EV1E)**. E10.5 *VavCre:loxGpr56* cKO AGM cells (CD41^lo^/CD45^+^; **Fig1E, Fig EV1F**) were similarly examined. A significant decrease in relative *Gpr56* mRNA was found in the cKO cells as compared to WT **(Fig1F)**. *Gpr56* deletion was verified by DNA-PCR **(Fig EV1G)**. Although there appears to be a trend towards reduced CFU-C numbers in CD41^lo^/CD45^+^ cKO AGM cells, it was not significant **(Fig1G)** and the percentages of phenotypic hematopoietic cells in the E10.5 cKO AGM were unchanged **(Fig1H)**. Together, these data indicate that loss of Gpr56 affects HPCs in the E9YS, but has little-to-no affect on E10.5 AGM hematopoiesis.

### HPC and HSC function is largely unaffected in E13.5 *Gpr56* cKO fetal liver

To examine whether Gpr56 loss affects definitive HSC and HPCs in FL, LSK-SLAM cells were isolated from E13.5 *VavCre:loxGpr56* and *VECCre:loxGpr56* cKO embryos(**Fig1I**). RT-PCR verified reduced levels of *Gpr56* transcripts in cKO cells as compared to WT (**Fig1J**) and DNA-PCR verified *Gpr56* deletion in *VavCre:loxGpr56* embryos (**Fig EV2A, upper**). The frequency of cKO HPCs (CFU-C/100 LSK-SLAM; **Fig1K**) and number of LSK-SLAM cells per E13.5 FL (**Fig1L**) in cKO lines were unchanged as compared to WT. When DNA from individual CFU-C were tested by *Gpr56*-PCR, 14/14 colonies from *VavCre* FL (**Fig EV2A**,**lower**), and 28/29 colonies from *VECCre* FL (**Fig EV2B**) had both *Gpr56* alleles recombined. Thus, *Gpr56* is dispensable for FL HPC growth and function.

E13.5 FL cKO LSK-SLAM cells were examined for HSC activity by *in vivo* transplantation (**Fig2A**). Adult irradiated recipients injected with one FL-equivalent of LSK-SLAM cells (∼100 HSCs(Kim, He et al., 2006)) from V*avCre:loxGpr56* cKO and WT donors showed no difference in the percentage of mice engrafted or peripheral blood (PB) donor-chimerism (4 and 16 weeks post-injection; **Fig2B**). Donor-chimerism in the bone marrow (BM), spleen (Sp), thymus (Thy) and lymph nodes (LN) of cKO recipients at 18 weeks (**Fig2C**) was equivalent to WT. Thus, Gpr56 appears dispensable for engraftment by large numbers of FL HSCs.

**Figure 2.**
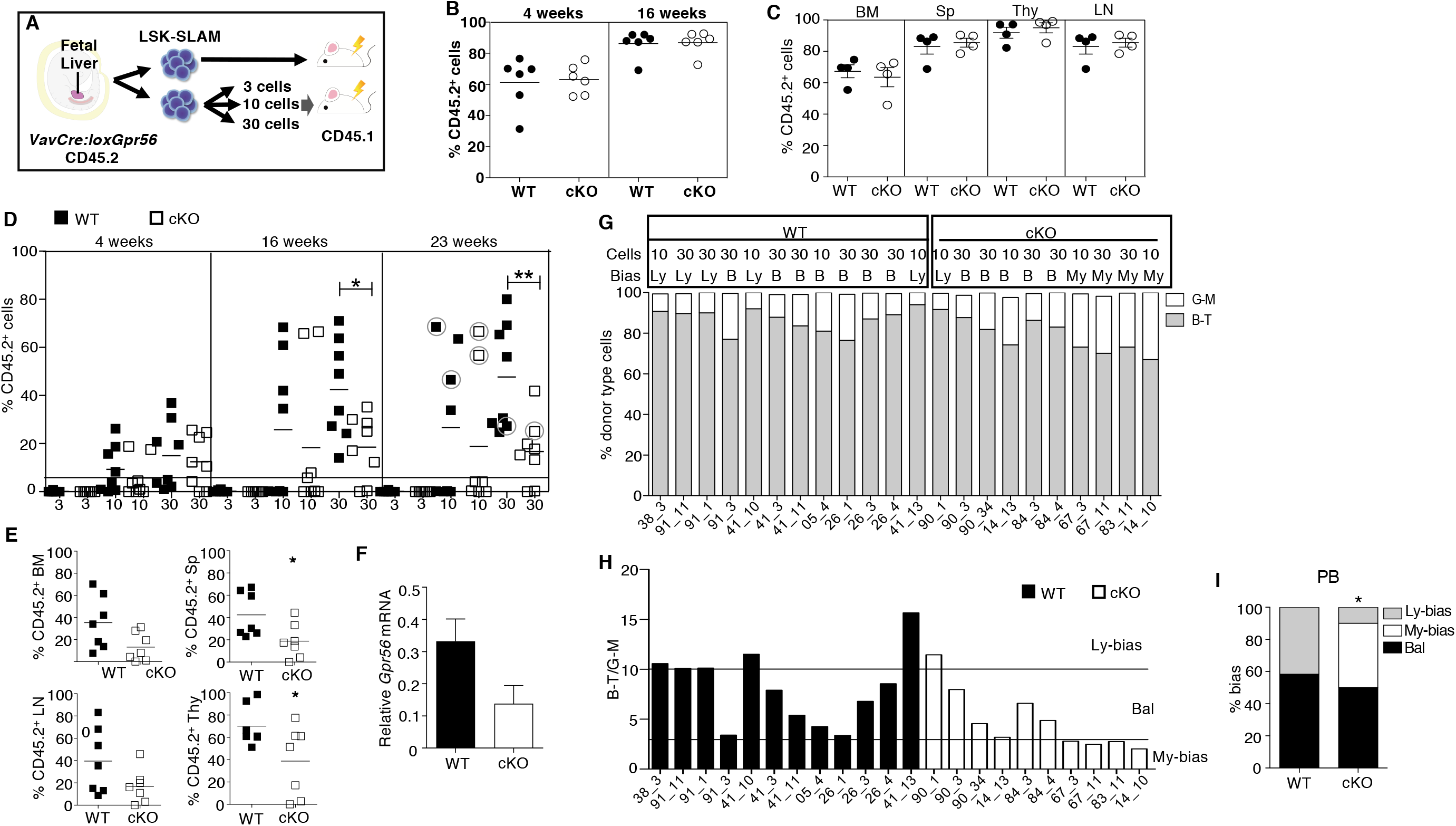
Clonal LSK-SLAM fetal liver cell *in vivo* transplantations reveal decreased engraftment levels and myeloid-lineage bias for Gpr56 deficient HSCs. **A)** Experimental setup for *in vivo* transplantation of E13.5 fetal liver (FL) sorted LSK-SLAM cells. **B and C)** One FL equivalent of LSK-SLAM sorted E13.5 cells from WT and *VavCre:loxGpr56* cKO embryos (Ly5.2) were injected into adult irradiated recipients (Ly5.1). Donor cell chimerism in recipient **B)** peripheral blood (PB) at week 4 and 16 post-transplant and **C)** hematopoietic tissues at week 18 post-transplant. Analysis is by Ly5.1/Ly5.2 flow cytometry. Mean±SEM are shown. n=6 per group. BM=bone marrow; Sp=spleen; Thy=thymus; LN=lymph nodes. **D)** Limiting dilution clonal HSC transplantation. Percentage of donor cell engraftment of individual adult irradiated recipient mice as measured by Ly5.1/Ly5.2 flow cytometry of PB at 4, 16 and 23 weeks post-injection of 3, 10 and 30 LSK-SLAM E13.5 FL cells. Wild type controls (WT=black) and *VavCre:loxGpr56* knockout (cKO=white). n=8 per group. Horizontal line at 5% indicates cutoff for reconstitution. Horizontal lines indicate average percentage engraftment. Circled individual symbols indicate mice used for secondary transplantations. *p≤0.05; **p≤0.01. **E)** Percentage of donor cell engraftment at 23 weeks post-injection as measured by Ly5.1/Ly5.2 flow cytometry of bone marrow (BM), spleen (Sp), lymph node (LN) and thymus (Thy) cells from 7 recipient mice injected with 30 LSK-SLAM FL cells. *p≤0.05. **F)** RT-PCR relative expression of *Gpr56* normalized to *β-actin* in LSK BM cells from primary recipients injected with WT and *VavCre:loxGpr56* cKO sorted E13.5 FL cells (n=5). Mean±SEM are shown. **G)** Percentage of lymphoid and myeloid cell contribution in PB of 22 individual adult irradiated recipient mice as measured by flow cytometry of peripheral blood at 23 weeks post-injection of 10 and 30 LSK-SLAM WT control and *VavCre:loxGpr56* cKO E13.5 FL cells. **H)** Ratio of B-T lymphoid and G-M myeloid for cohort of recipients in panel G with indicators for limits of lymphoid-biased, balanced and myeloid-biased HSC outputs. Ly=lymphoid bias; B=balanced; My=myeloid bias. >87% B-T=Ly; 75-87% B-T=B; <75% B-T=My. **I)** Percentages of lymphoid-biased, balanced and myeloid-biased HSC engrafted recipients from panel H. Fisher exact test determined statistically significant differences in the My-bias fraction. *=p≤0.05.

### Clonal transplantation reveals a role for Gpr56 in FL HSC lineage-bias

To test if Gpr56 affects HSC quality in clonal *in vivo* transplantation, 3, 10 and 30 LSK-SLAM FL cells from E13.5 *VavCre:loxGpr56* embryos and WT littermates were injected into irradiated adult mice (**Fig2D**). Injection of 3 LSK-SLAM cells failed to reconstitute. Injection of 10 or 30 WT cells led to long-term (23 week) reconstitution (≥5% donor-chimerism) in 75% (12/16) of recipients, whereas only 50% (8/16) of recipients receiving cKO cells showed reconstitution. At both 16 and 23 weeks post-injection of 30 cells, the average percentage PB donor-chimerism in cKO recipients was significantly decreased as compared to WT. Also, the percentage donor-chimerism was decreased in the BM, Sp, LN and Thy of cKO recipients (**Fig2E**). *Gpr56* downregulated expression was verified in CD45.2^+^ LSK cells from the BM (**Fig2F**), thus confirming Gpr56-related HSC dysfunction.

The donor-derived lymphoid-myeloid ratio(Sieburg, Cho et al., 2006) was examined in the PB of cKO and WT E13.5 LSK-SLAM FL reconstituted recipients at 23 weeks (**Fig2G**). Lineage output was determined by B plus T lymphoid cell percentages to granulocyte plus macrophage percentages (B+T^value^/G+M^value^), with ratios of >10, 10-3 and 3-0 considered lymphoid-biased, balanced or myeloid-biased respectively (**Fig2H**). The 12 WT HSC recipients were divided between lymphoid-biased (5) and balanced (7) output. In contrast, the 10 cKO HSC recipients showed 1 lymphoid-biased, 5 balanced and 4 myeloid-biased **(Fig2H)**. The increase in myeloid-biased HSCs was significant (**Fig2I**). The BM cell lineage output of some of this cohort (**Fig EV2C, D**) showed a similar but not significant myeloid-bias in cKO HSC recipients. Secondary transplantation recipients of primary BM Ly5.2 cells showed 0/12 cKO recipients repopulated, whereas 4/12 WT recipients had PB-chimerism of 2.5-15.3% (**Fig EV2E**). These data suggest that Gpr56 maintains the *in vivo* quality of HSCs, preserving lymphoid-biased and balanced HSC lineage output.

### Gpr56 is expressed in ESC-derived hematopoietic cells

To further examine the role of Gpr56 during development/differentiation, we used *Gata2Venus* (*G2V*) mouse reporter ESCs(Kaimakis, de Pater et al., 2016), which facilitate the isolation of HS/PCs in the Venus-expressing (V^+^) cell fraction for molecular and functional analyses(Kaimakis et al., 2016, Kauts, Rodriguez-Seoane et al., 2018) (**Fig EV3**, gating and FMOs). *G2V* ESCs were differentiated and harvested at several time points (**Fig3A**). RT-PCR analysis on cells from day (d)0, 6, 10 and 12 cultures showed significant increases in *Gpr56* transcript levels, peaking at d10 and d12 in unsorted (**Fig EV4A**) and V^+^ cells (**Fig EV4B**). Low level of *Gpr56* expression was found in non-hematopoietic V^-^ cells as expected(Ackerman et al., 2015). Western blotting of whole cell extracts from unsorted differentiated *G2V* ESCs showed high level Gpr56 protein expression at d12 as compared to d0 (**Fig EV4C, left**) and in d12 V^+^ as compared to V^-^ cells (**Fig EV4C, right**). Thus, *G2V* ESCs are a suitable platform to examine the role of Gpr56 during *in vitro* hematopoietic differentiation.

**Figure 3.**
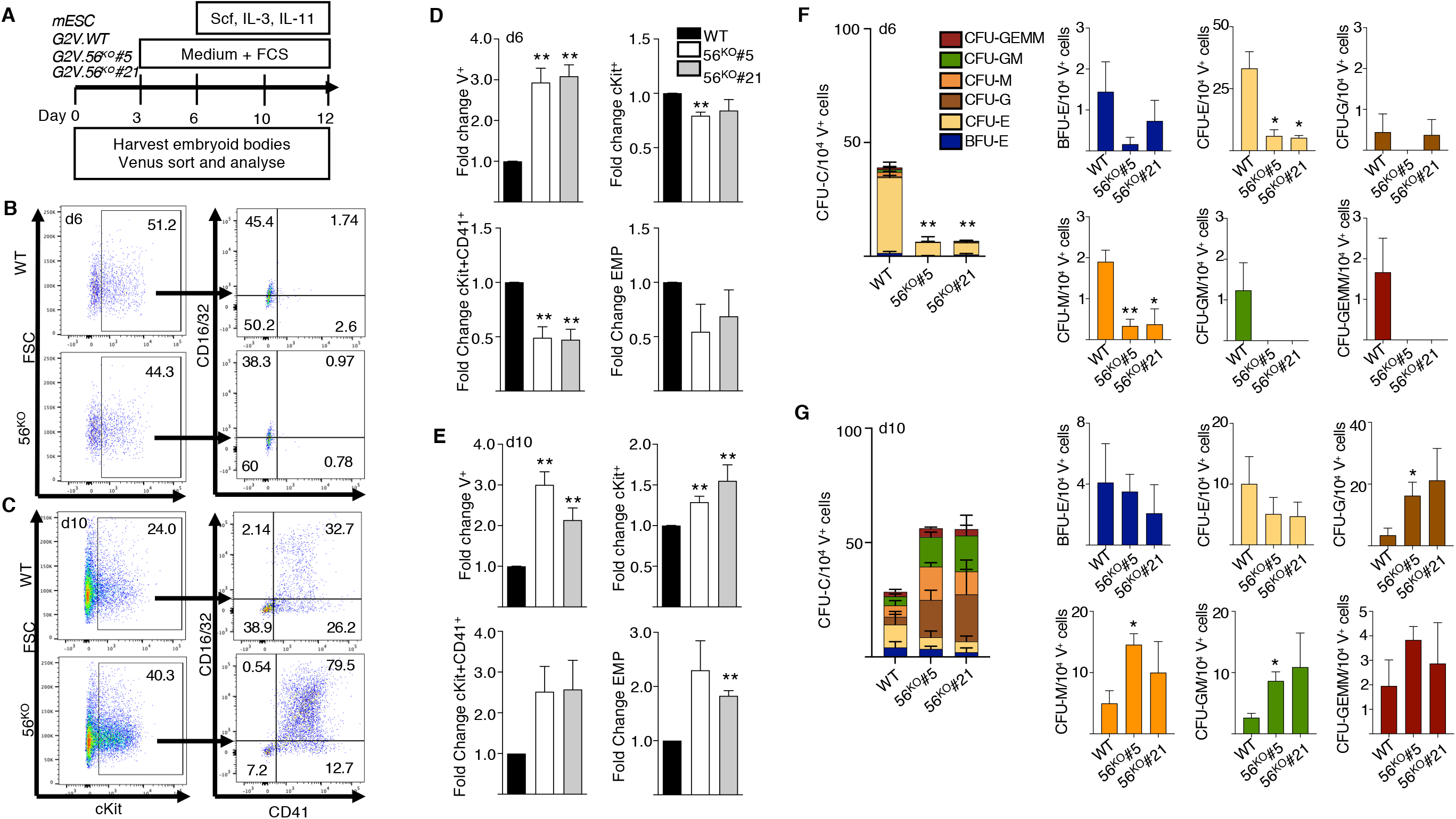
Gpr56 affects hematopoietic output *in vitro*. **A)** ESC differentiation culture methodology. Mouse *Gata2Venus (mG2V)* ESC (wild type (WT) and *56*^*KO*^) were differentiated in hematopoietic factor-containing medium for several days and unsorted/sorted Venus^+^ (V^+^) and Venus^-^ (V^-^) cells were examined for *Gpr56* mRNA (RT-PCR) and Gpr56 protein (Western blot; WB) expression. Representative flow cytometric plots of **B)** day 6 and **C)** day 10 *G2V.WT* and *G2V. 56*^*KO*^ differentiation cultures showing percentages of CD16/32 and CD41 cells in Venus+cKit^+^ gate. CD41^+^cKit^+^ CD16/32^+^ = EMP (erythro-myeloid progenitors). Fold change in the percentages (Mean±SEM) of Venus^+^, cKit^+^, cKit^+^CD41^+^ and EMPs in **D)** day 6 and **E)** day 10 *G2V.WT* and *G2V. 56*^*KO*^ (Clones #5 and #21) differentiation cultures. n=3. Hematopoietic potential of *G2V.WT* and *G2V.Gpr56* ^*KO*^ (Clones #5 and #21) HPCs was determined at day 6 **F)** day 6 and **G)** day 10 of differentiation by colony forming unit-culture (CFU-C) assay. CFU-C per 10,000 V^+^ plated cells is shown. Distinct colony types are indicated by color. n=3; Mean±SEM; CFU-GEMM=granulocyte, erythroid, megakaryocyte, macrophage; -GM=granulocyte, macrophage; -G=granulocyte; M=macrophage; E=erythroid; BFU-E=burst forming unit-erythroid.

### *Gpr56* knockout affects hematopoietic output during ESC differentiation

*Gpr56* knockout *G2V* ESC lines (*56*^*KO*^) were generated by *CRISPR/Cas9* editing. Guide RNAs (*gRNA*) to exon 2 (common to all transcript variants (**Fig EV5A**) were cloned (*pSP-Cas9-2A-GFP)* and transfected into *G2V* ESCs (**Fig EV5B**). At 48 hours post-transfection, ∼10% of *G2V* ESCs were GFP^+^ (**Fig EV5C, D**). Day 6 differentiated *G2V* clones (#5, #21) showed a complete lack of Gpr56 protein (**Fig EV4D**) as compared to WT (#1, #2). The total absence of *Gpr56* transcripts in clone #5 as compared to WT ESC was verified during time-course *in vitro* differentiation (**Fig EV4E**), thus confirming the successful knockout of *Gpr56*.

Phenotypic HPC(McGrath, Frame et al., 2015) and CFU-C analyses were performed on cells from d6 (**Fig3B, D, F)** and d10 (**Fig3C, E, G**) *56*^*KO*^ and WT ESC cultures. The percentage of V^+^ cells in both the d6 and d10 *56*^*KO*^ (#5 and #21) cultures was significantly increased (2.8±0.95 fold) as compared to WT (**Fig3D, E**). Despite this increase, at d6 a significant decrease in the percentage of cKit^+^ (#5) and cKit^+^CD41^+^ cells in the V^+^ fraction (#5 and #21) of *56*^*KO*^ cultures was found compared to WT (**Fig3D**). As expected EMPs are not detected (Kauts, Rodriguez-Seoane et al., 2018). In contrast d10 *56*^*KO*^ ESC cultures showed a significant increase in cKit^+^ (#5 and #21; 1.3±0.2) and phenotypic EMP percentages (#21; 1.82±0.1) (**Fig3E**).

When d6 V^+^ cells were tested for HPC function (**Fig3F**), both *56*^*KO*^ clones were significantly reduced (4-fold) in CFU-C/10^4^ V^+^ cells as compared to WT, indicating an almost complete absence of hematopoietic progenitors. The significant decrease was in CFU-E and CFU-M. In contrast, V^+^ cells from d10 *56*^*KO*^ (#5, #21) cultures (**Fig3G**) revealed a trend (p=0.33, p=0.68) towards increased CFU-C/10^4^ cells as compared to *WT* V^+^ cells. Significant increases in CFU-G, -M and -GM frequencies for clone #5 were observed. Hence, the reduction in d6 early HPCs and the contrasting increase in myeloid lineage progenitors at d10 of *56*^*KO*^ ESC differentiation agrees with *in vivo Gpr56* cKO results and suggests that Gpr56 plays distinct roles during hematopoietic development/differentiation.

In the developing brain, Col3a1 is a Gpr56 ligand(Giera, Deng et al., 2015, Luo, Jeong et al., 2011) and Gpr56 signals intracellularly through Rho/ROCK(Ackerman et al., 2015). To test these routes of activation in hematopoietic cells, Col3a1 or ROCK inhibitor Y-27632 were added to *in vitro* differentiating *G2V* ESCs. Cultures (d12) showed no changes in the number of ESC-derived V^+^ hematopoietic cells for either condition (**Fig4A**). When HPC activity was analyzed, a trend towards reduced total CFU-C numbers was found with the ROCK inhibitor, whereas an increased trend was found with Col3a1 (**Fig4B**). Examination of specific colony types revealed a significant increase in CFU-M and CFU-GEMM in the presence of Col3a1 (**Fig4C**). The ROCK inhibitor decreased BFU-E, CFU-M and CFU-GM (**Fig4C**). Our single cell RNAseq dataset(Vink, Calero-Nieto et al., 2020) of HSC, HPCs and endothelial cells in E10.5/11.5 intra-aortic cluster (CD31^+^G2V^med^cKit^hi^) was examined for *Col3a1, Gpr56* and *Arrb2*(Kroeze, Sassano et al., 2015, Lefkowitz & Shenoy, 2005) (effector of Gpr56 activation) (**Fig4D**). *Gpr56* was found highly expressed in the hematopoietic cluster (HC1) whereas *Col3a1* was highly expressed in the endothelial-like cluster (EC). *Arrb2* was expressed at varying levels in both. These data agree with published expression analysis of AGM hematopietic and niche populations(Pimanda & Gottgens, 2010) and support the notion that Col3a1 is a likely ligand for Gpr56 in hematopoietic development.

**Figure 4.**
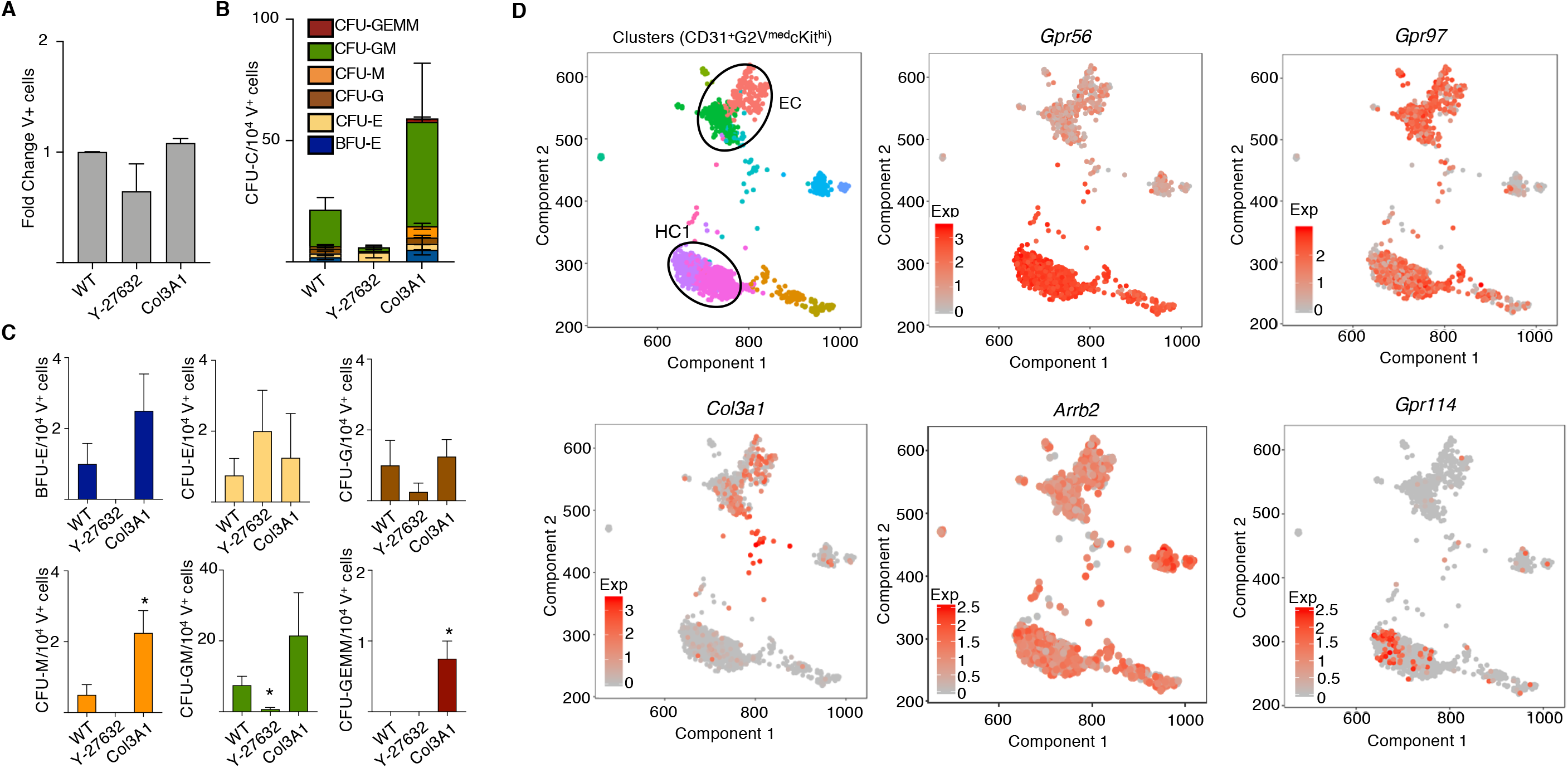
Functional and gene expression analyses of Gpr56 downstream effectors. **A)** Frequency of Venus+ cells and **B, C)** CFU-C colonies of d12 *G2V.WT* sorted cells treated with ROCK inhibitor (Y-27632) and collagen (Col3a1). **D)** Spring analysis of AGM-derived HSPCs(Vink et al., 2020) displays the differential expression of *Gpr56*, Collagen (*Col3a1*), beta arrestin (*Arrb2*), *Gpr97* and *Gpr114* in the two cell clusters (EC and HC1) of highly enriched HSPCs (CD31^+^G2V^med^cKit^hi^) from E10.5/11.5 AGM IAHC cells.

### Mouse *Gpr97* rescues HS/PC generation in Zebrafish *gpr56* morphants

The discrepancy between the involvement of *Gpr56* in zebrafish and mouse HS/PC development may be a result of GPCR redundancy(Solaimani Kartalaei et al., 2015). Three closely-related Gpr genes (5’ *Gpr114;Gpr56;Gpr97* 3’) are contained within the 122Kb mouse locus. In contrast, zebrafish *gpr56* and *gpr97* are separated by 36Mb (**Fig5A**) and no *gpr114* exists. Since mouse *Gpr97* and *Gpr114* are highly-homologous to *Gpr56* and co-expressed in enriched AGM HS/PC populations(Solaimani Kartalaei et al., 2015), they may compensate for *Gpr56* loss. To test for functional redundancy, mouse *Gpr97* and *Gpr114* mRNA (as shown for mouse *Gpr56(Solaimani Kartalaei et al., 2015)*) were injected into zebrafish embryos along with *gpr56* morpholino (MO) (**Fig5B**). Morphant *gpr56* embryos co-injected with mouse *Gpr97* mRNA showed hematopoietic rescue (*myb in situ* hybridization at 24-30hpf), whereas mouse *Gpr114* mRNA did not. Additionally, transgenic zebrafish embryos (*CD41GFP*:*Flt1RFP* report HS/PC generation) were injected with the *gpr56* MO alone or in combination with zebrafish *gpr56* or mouse *Gpr56, Gpr97* or *Gpr114* mRNA. Double expressing CD41^dim^Flt1^+^ (GFP^+^RFP^+^) HS/PCs in the caudal hematopoietic cell tissue (CHT) at 48 hpf were counted (**Fig5C, D**). Significant increases in the average number of HS/PCs were found in the morphants injected with zebrafish *gpr56*, mouse *Gpr56* and mouse *Gpr97* mRNA versus control morphants, and HS/PC numbers were equivalent to those in WT controls. In contrast, mouse *Gpr114* mRNA did not restore production of HS/PC. Alignment comparisons revealed a high degree of homology between mouse Gpr97 and both zebrafish gpr56 and mouse Gpr56 in all 4 protein domains (**Fig5E**), thus suggesting that Gpr97 may function redundantly in the mouse embryos in the absence of Gpr56.

**Figure 5.**
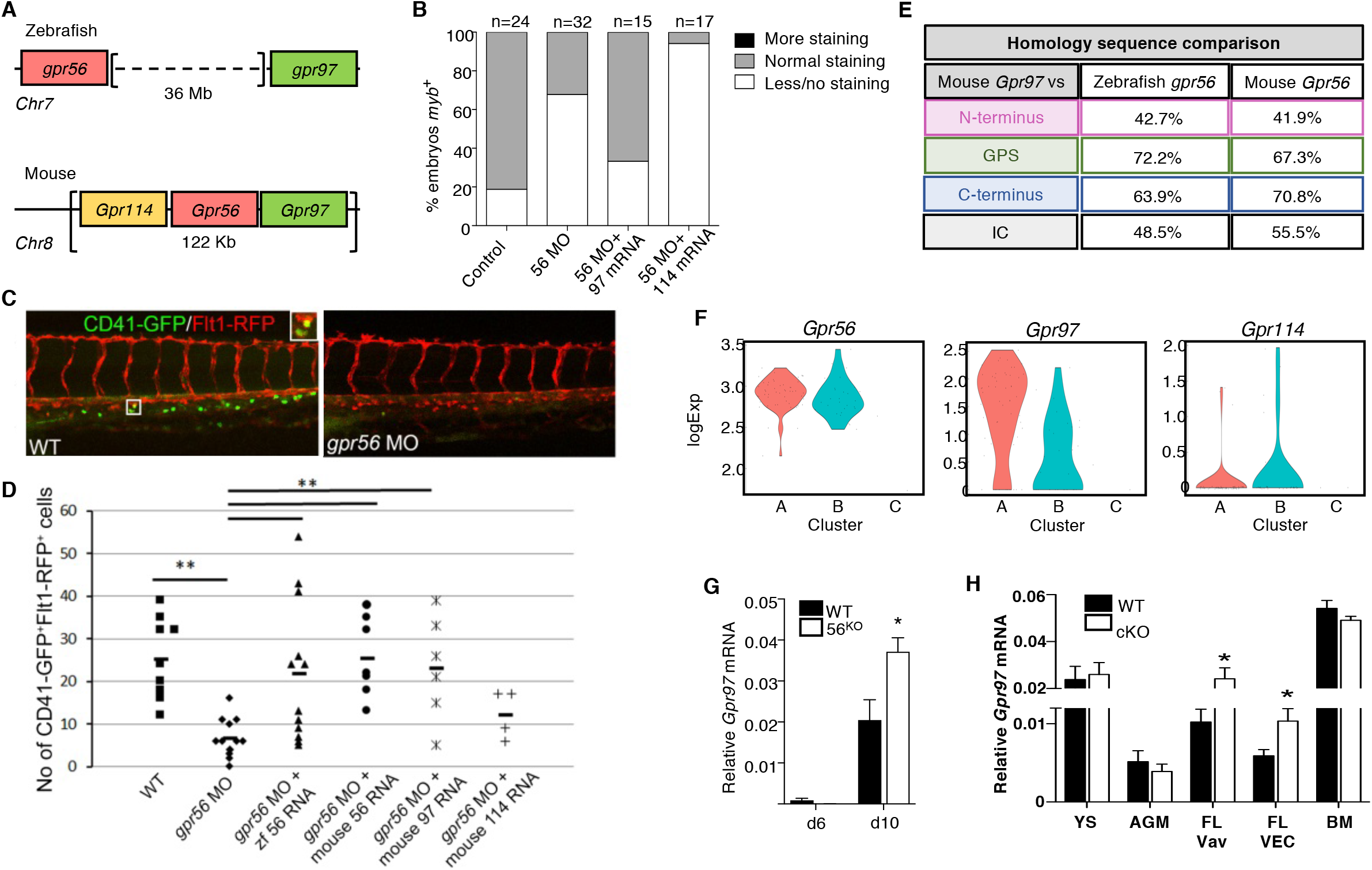
Redundant function of mouse Gpr97 in *Gpr56* morphant zebrafish and *Gpr97* expression in mouse ESCs and embryonic tissues. **A)** Schematic of the *Gpr56* locus in zebrafish and mouse. Two highly homologous genes, *Gpr114* and *Gpr97* are located 5’ and 3’ respectively within the mouse *Gpr56* locus. **B)** *In situ* hybridization with the HSC marker *myb* at 30 hpf. Quantification of zebrafish embryos injected with *gpr56* morpholino +/- *gpr97* or *gpr114* mRNA. n= the number of embryos examined per condition. **C)** Representative images of CD41GFP^dim^Flt1RFP^+^ cells (yellow fluorescence) in the caudal hematopoietic tissue (CHT) of *gpr56* morpholino (MO) injected and control (WT) double transgenic zebrafish embryos at 48 hpf. CD41=green; Flt1= red; double positive definitive HS/PC=yellow (left panel). **D)** Rescue of HS/PC production as determined by the number of CD41GFP^dim^Flt1RFP^+^ cells in the CHT of WT and *gpr56* morphant zebrafish at 48 hpf when zebrafish *gpr56*, mouse *Gpr56*, mouse *Gpr97* and mouse *Gpr114* mRNA was injected. n=2 (9, 12, 12, 7, 6, 4 embryos injected and analysed, respectively); to change? horizontal bars=mean; **p<0.01. **E)** Percentages of amino acid sequence homology of the four domains of mouse Gpr97 versus zebrafish gpr56 and mouse Gpr56. **F)** Violin plots showing logExp of *Gpr56, Gpr97 and Gpr114* in the 85 CD27^+^ cells within HC1 (Fig4F) generated from Vink et al., 2020 single cell RNA database. HC1 CD27+ cells are segregated into subclusters A, B, C. **G)** RT-PCR analysis of relative *Gpr97* expression (normalized to *β-actin*) in d6 and d10 diferentiated *G2V* WT and *G2V.56*^*KO*^ ESCs from hematopoietic differentiation cultures (same V^+^cKit^+^ samples as shown in Fig EV4B for relative *Gpr56* expression). n=3. Mean±SEM. **H)** RT-PCR analysis of relative *Gpr97* expression in E9 *VECCre:loxGpr56* cKO yolk sac (YS) cells (n=3), E10.5 *VavCre:loxGpr56* AGM cells (n=3), *VavCre:loxGpr56* and *VECCre:loxGpr56* cKO fetal liver (FL) cells (n=5 and n=8, respectively) and LSK bone marrow cells (n=5) from primary recipients of *VavCre:loxGpr56* cKO cells. *p≤0.05.

### *Gpr97* expression is upregulated in the absence of *Gpr56 in vivo* and *in vitro*

Mouse *G2V*.*WT* ESCs were examined by RT-PCR for *Gpr97* expression during *in vitro* hematopoietic differentiation. *Gpr97* expression increased significantly from d6 to d12 in the V^+^ fraction (**Fig EV4F**). Violin plots generated from our scRNAseq datasets of enriched AGM HS/PCs(Vink et al., 2020) (**Fig5F**) show high expression of *Gpr56* in the 85 CD27^+^ cells in HC1. These cells separate into three subclusters (A, B, C). Cluster B contains all the HSC activity(Vink et al., 2020). As compared to *Gpr56* logExp, *Gpr97* transcripts are 10-fold lower in A and 100-fold lower in B, and *Gpr114* transcripts are barely detectable. The individual cells in the Spring analysis (**Fig4D**) shows co-expression of *Gpr97* in many *Gpr56* expressing cells in HC1, with co-expression of *Gpr114* in very few. Thus, low *Gpr97* expression overlaps with high *Gpr56* expression in some enriched HS/PCs in mouse IAHCs.

*Gpr97* levels were examined in i*n vitro* differentiated *Gpr56*^*KO*^ ESCs (**Fig5G**) and *in vivo VECCre:loxGpr56* and *VavCre:loxGpr56* cKO YS, AGM FL and BM hematopoietic cells (**Fig5H**). When compared to differentiated WT G2V.ESCs, significantly increased expression of *Gpr97* expression was found in the d10 (but not d6) V^+^ fraction. E9 YS CD31^+^ cells and E10.5 CD41^lo^/CD45^+^ AGM cells showed similar levels of *Gpr97* expression in WT and *Gpr56* cKO cells. In contrast, E13.5 FL LSK-SLAM cells showed a significant upregulated expression of *Gpr97* in *Gpr56* cKO (∼1.5-fold) versus *WT* cells. The donor-derived LSK BM cells from transplant recipients of 10 or 30 cKO E13.5 LSK-SLAM FL cells showed comparable levels of relative *Gpr97* expression to WT LSK BM cells. Thus, Gpr97 may function redundantly in the definitive stage of hematopoietic development both *in vitro* (d10) and *in vivo* (FL), but not in early development or in BM of transplant recipients.

### Deletion of both *Gpr56* and *Gpr97* impairs ESC-derived hematopoiesis

To explore the functional contribution of Gpr97 during *in vitro* differentiation upon loss of *Gpr56*, we generated a mouse *G2V*.*56*^*KO*^*97*^*KO*^ ESC line (**Fig6A, Fig EV6**). *CRISPR/Cas9* editing with gRNAs targeting *Gpr97* exons 2 and 10 was used to knockout all potential isoforms and/or decay RNA transcripts. Clone B3 (*56*^*KO*^*97*^*KO*^) was used for further studies. The absence of *Gpr97* transcripts in B3 (d6 and d10) cells was confirmed by RT-PCR (**Fig6B**). Hematopoietic output of d10 *56*^*KO*^*97*^*KO*^, *56*^*KO*^ and *WT* ESC was examined by flow cytometry (**Fig6C**). The same high percentage of viable cells (>90%) was found for all three ESC lines (**Fig6D**) and whereas the fold-increase of V^+^ cells in the *56*^*KO*^ cultures was significant, *56*^*KO*^*97*^*KO*^ cultures showed no significant changes in V^+^ cells as compared to WT. However, significant fold-decreases in CD45^+^ cells and CD41^+^cKit^+^CD16/32^+^ (EMP) cells were observed in the *56*^*KO*^*97*^*KO*^ differentiated cells as compared to WT. This suggests that both Gpr56 and Gpr97 are involved in HPC cell generation and/or differentiation.

**Figure 6.**
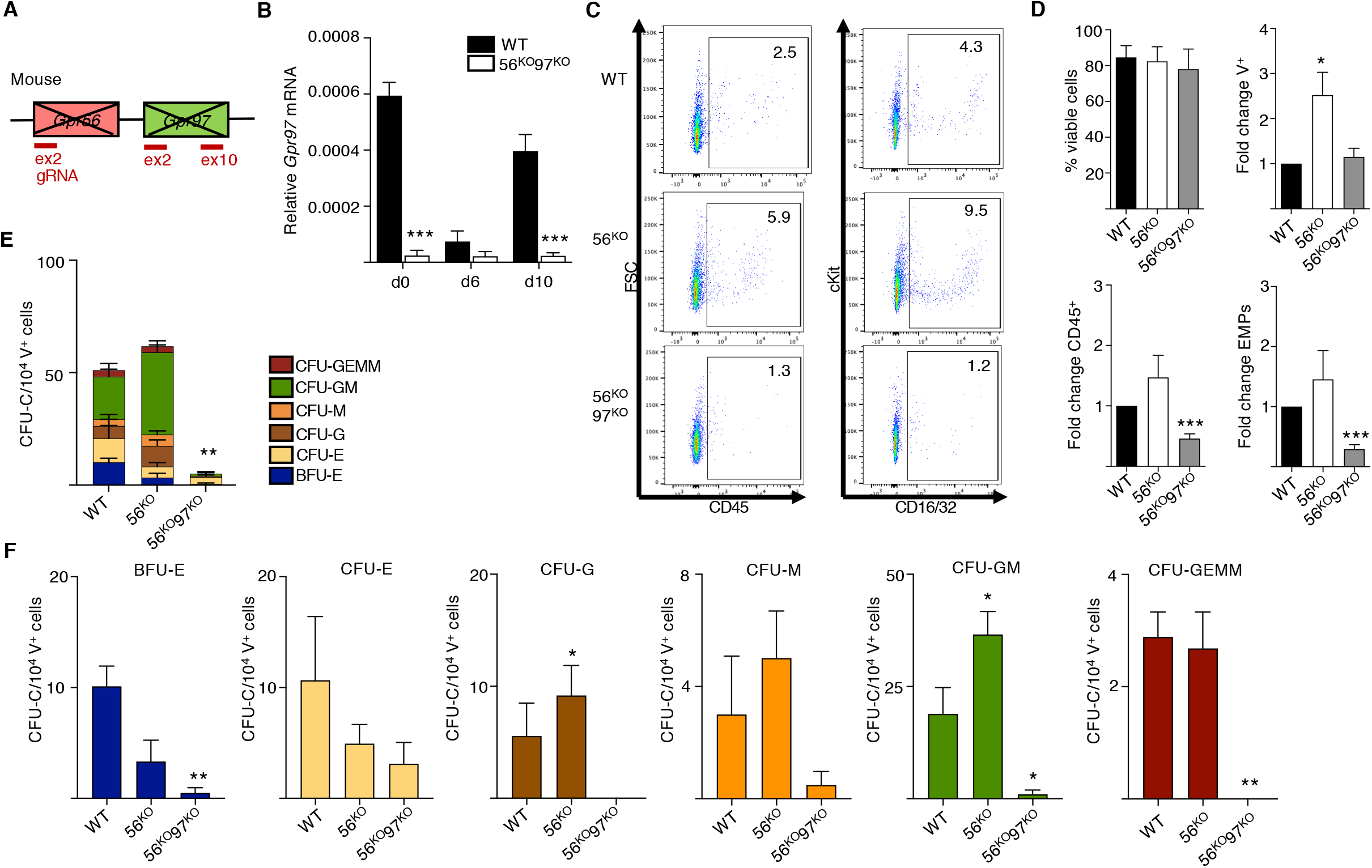
Decreased production of HPCs in the absence of *Gpr56* and *Gpr97*. **A)** Schematic of *Gpr56* and *Gpr97* deletion in mouse *Gpr56* locus. **B)** RT-PCR analysis of time course hematopoietic differentiation cultures of *G2V* WT and *G2V.56*^*KO*^*97*^*KO*^ ESCs. Relative *Gpr97* expression (normalized to *β-actin*). **C)** Representative flow cytometry plots for CD45^+^ hematopoietic cells and EMPs (CD41^+^cKit^+^CD16/32^+^) from d10 differentiated *G2V* WT, *G2V.56*^*KO*^ and *G2V.56*^*KO*^*97*^*KO*^ ESC cultures. Percentages shown in each quadrant. **D)** Percentage of viable cells and fold-change in percentages of Venus(V)^+^, CD45^+^, and EMPs from day10 differentiated *G2V* WT, *G2V.56*^*KO*^ and *G2V.56*^*KO*^*97*^*KO*^ ESC cultures. n=3. Mean±SEM. **E)** Frequency of colony forming unit-culture cells (CFU-C) in day 10 differentiated *G2V* WT, *G2V.56*^*KO*^ and *G2V.56*^*KO*^*97*^*KO*^ ESC cultures. CFU-C per 10^4^ Venus^+^ cells is shown with colony types indicated by color. CFU-GEMM=granulocyte, erythroid, megakaryocyte, macrophage; - GM=granulocyte, macrophage; -G=granulocyte; M=macrophage; E=erythroid; BFU-E=burst forming unit-erythroid. n=3. Mean±SEM. **F)** Output of CFU-C per 10^4^ Venus^+^ in the different colony types: BFU-erythroid, CFU-E=erythoid, -G=granulocytes, - M=macrophage, -GM=granulocytes, macrophage, -GEMM=multilineage.*p≤0.05; **p≤0.01; ***p≤0.001.

CFU-C assays were performed to quantitate HPC production in the V^+^ fraction of d10 56^*KO*^*97*^*KO*^, *56*^*KO*^ and *WT* ESCs. In contrast to *56*^*KO*^ V^+^ cells in which total CFU-C frequency was similar or slightly increased to *WT* V^+^ cells, *56*^*KO*^*97*^*KO*^ V^+^ cells showed a significant large reduction in CFU-C (**Fig6E**). Moreover, there was a significant reduction in the frequency of *56*^*KO*^*97*^*KO*^ derived BFU-E, CFU-GM and CFU-GEMM (**Fig6F**) as compared to *WT*. Thus, we conclude that definitive HPC production is dependent upon expression of *Gpr56* and *Gpr97*.

## Discussion

We have shown that two closely-related G-protein coupled receptors, Gpr56 and Gpr97, function in the production and qualitative output of HS/PCs during mouse embryonic development. While the knockout of *Gpr56* alone in mouse ESCs partially affects HPC production and biases output to the myeloid lineage, the double knockout of *Gpr56* and *Gpr97* severely reduces definitive HPC production and differentiation. Moreover, *Gpr56* deleted FL HSCs show myeloid lineage-bias and decreased robustness *in vivo*. Together, these data support a role for *Gpr56* in enforcing multilineage potential on HSCs in the embryo, reveal a developmenatal stage-dependent redundant function for Gpr97, and suggest that *Gpr97* upregulation promotes the myeloid-bias of HS/PCs.

### *Gpr56* deletion provides new insights into the regulation of mammalian hematopoiesis

Despite the high expression of Gpr56 in quiescent adult BM HSCs(Rao et al., 2015), the function of this adhesion-GPCR in HSCs has been controversial. Whereas a study in human leukemic cell lines and a germline *Gpr56*^*-/-*^ mouse model claimed that EVI1-regulated *GPR56* maintains the HSC pool in BM niches(Saito et al., 2013), others reported that this same germline knockout mouse was not impaired in BM HS/PC maintenance or function during homeostasis or stress(Rao et al., 2015). This was suggested to be due to a hypomorphic *Gpr56* allele (low expression of S4 splice variant) or to the mouse background. Our finding of upregulated *Gpr56* expression in mouse embryo IAHC cells and the impairment of aortic HS/PC generation in *Gpr56* morphant zebrafish embryos(Solaimani Kartalaei et al., 2015) prompted re-examination of Gpr56 function during mouse hematopoietic development.

In our studies, *Gpr56* was conditionally-deleted (exons 4,5 and 6 floxed allele(Giera et al., 2015)) in HPCs and HSCs of embryos with *VECCre* and *VavCre*. E9 YS HPC numbers were reduced, but E13.5 FL LSK-SLAM HSC numbers were unaffected. However, clonal *in vivo* transplantations of FL LSK-SLAM cells revealed qualitative changes in HSCs. *Gpr56-*loss affected the lineage-bias of FL HSCs. *In vitro Gpr56*^*KO*^ ESC hematopoietic differentiation cultures at d6 were decreased in phenotypic and functional HPCs, similar to E9 cKO YS. However, d10 *Gpr56*^*KO*^ ESCs were slightly increased in phenotypic and functional HPCs and, like E13.5 cKO FL HSCs, they showed significant myeloid-bias compared to the control. Hence, Gpr56 likely affects early embryonic hematopoietic generation quantitatively, and at later developmental times and/or in distinct microenvironments it has a role in HSC quality.

### Gpr97 influences hematopoietic development in the absence of Gpr56

Compared to the requirement of *Gpr56* for HS/PC generation in zebrafish embryos, the rather minor roles for *Gpr56* in mouse hematopoietic development remained puzzling. If Gpr56 was essential for mouse HS/PC generation, embryonic lethality would be expected. Instead, homozygous *Gpr56* mutant adults could be produced, and the characteristics of mutant BM HS/PCs appeared minimal(Rao et al., 2015, Saito et al., 2013). Our rescue of *Gpr56* morphant zebrafish HS/PCs with mouse *Gpr56* and *Gpr97*, but not *Gpr114* mRNA raised the possibility of receptor redundancy. The linkage of *Gpr56* and *Gpr97* and the co-expression of these highly homologous receptors in the mouse, together with the loss of genomic synteny in zebrafish allowed us to identify this redundancy. Importantly, *gpr97* is not expressed in the zebrafish embryos at the time of HPSC generation(Harty, Krishnan et al., 2015).

Similar to mice, the human *GPR56* locus contains highly-homologous *GPR114* and *GPR97* genes in the same 5’ to 3’ configuration. The evolutionary conservation of mouse and human genes/proteins, together with similarities in the embryonic development of mouse and human hematopoietic systems, suggests an overlap in the functions of Gpr56 and Gpr97 (explaining the minor effects seen in the mouse Gpr56 knockout models). Redundancy is supported by our results showing *Gpr97* expression in cells of E9 YS, E13.5 FL and adult BM, and also in cells of d6 and d10 ESC differentiation cultures. Importantly, in the absence of *Gpr56, Gpr97* expression was upregulated in E13.5 FL LSK SLAM cells and d10 differentiated ESC derived V^+^cKit^+^ hematopoietic cells. No changes were found in *Gpr97* expression in *Gpr56* cKO E9 YS, E9 AGM or adult BM. Thus, only in the definitive HSC stage in the embryo the loss of Gpr56 function is compensated by expression of Gpr97.

### Positive role of Gpr97 or negative role of Gpr56 for myeloid-bias?

Myeloid-bias was observed in cells from *in vitro* differentiated *Gpr56*^*KO*^ ESC cultures and *in vivo* in the recipients of *Gpr56* cKO FL HSCs. Previously, lineage-biased output of HSCs has been reported as an age-related characteristic(Benz, Copley et al., 2012, Cho, Sieburg et al., 2008, Geiger, de Haan et al., 2013, Verovskaya, Broekhuis et al., 2013). As measured by long-term *in vivo* clonal transplantation, BM HSCs from young mice yield lymphoid-biased and balanced output. BM from aged mice show a higher frequency of myeloid-biased HSC and a decreased self-renewal ability. The myeloid-bias of *Gpr56* cKO FL HSCs and lower-level of donor chimerism in primary recipients support a qualitative role(s) for Gpr56 in HSCs. We suggest that Gpr56 is responsible for maintaining the multipotency and robustness of HSCs throughout development. Indeed, these critical qualitative properties appear for the first time in midgestation mouse embryos when the first adult-repopulating HSCs are generated(Dzierzak & Bigas, 2018, Dzierzak & Speck, 2008). Highly-upregulated *Gpr56* expression localizes to emerging E10.5/11.5 IAHC cells at after they transition from endothelial cells(Solaimani Kartalaei et al., 2015) and we show that co-expression of *Gpr97* and *Gpr56* in single cell transcriptomes is highly enriched in AGM HS/PCs.

Because of overlapping expression, it remains unclear whether Gpr56 functions to retain multipotency or represses myeloid-bias, or whether Gpr97 (especially its upregulated expression in the absence of Gpr56) activates myeloid-bias. In human NK cells, GPR56 negatively regulates immediate effector functions(Chang, Hsiao et al., 2016). Human polymorphonuclear cells highly express GPR97, whereas monocytes and lymphocytes do not(Hsiao, Chu et al., 2018). These results highlight a likely balance of these GPCRs in specific subsets of hematopoietic cells as an important factor impacting cellular functions. As we have demonstrated here that ESC-derived hematopoiesis is severely reduced in the absence of both *Gpr56* and *Gpr97*, the specific functions of these two closely-related receptors can only be addressed through the generation of a double-knockout mouse or human cell line model. Our attempts to generate a *Gpr56*^*-/-*^*Gpr97*^*-/-*^ germline mouse have been unsuccessful, likely due to embryonic lethality. Future studies aim to understand the individual roles of these receptors during HSC development by generating a cKO model across both genes and comparing effects with single cKOs and/or individual rescue of *Gpr56* and *Gpr97*.

## Materials and methods

### Mice

*Gpr56fl*(Giera et al., 2015), *VavCre*(Stadtfeld & Graf, 2005) and *VECCre*(Chen, Yokomizo et al., 2009) mice were maintained and embryos generated by mating 2-6 month old *Gpr56fl*(C57BL/6) and *VavCre:loxGpr56* or *VECCre:loxGpr56* (C57BL/6HsdJOla) mice. Vaginal plug discovery was embryonic day(E)0. Staging was by somite counts. Ly5.1 mice (B6.SJL-*Ptprc*^*a*^*Pepc*^*b*^/BoyCrl, 2-4 months) were transplant recipients. Mice were maintained in University of Edinburgh animal facilities in compliance with a Home Office UK Project License.

### Flow cytometry

Yolk sacs (YS) and AGMs were digested (37°C, 45’) in 0.125% collagenaseT1(C0130, Sigma-Aldrich) in PBS+10%FBS. FL cells were dispersed (5x) through a 30G needle. YS and AGM cells were stained with CD41-eFluor450, CD31-BV605, CD16/32-PE, c-Kit-APC and CD45-AF450 and FL with LSK-SLAM markers (CD3^-^B220^-^Gr1^-^Ter119^-^ NK1.1^-^CD48^-^, Sca1^+^cKit^+^CD150^+^) (**Table EV1**) and analysed (LSR Fortessa/FlowJov10, BD).

EBs were PBS-washed, dissociated (37°C, 5-10’) in 500μl TrypLE Express (Gibco) and anti-CD41, CD45, CD16/32, and cKit stained (4°C, 30’). Dead cell exclusion was by Hoechst 33342 (Invitrogen) and gates set with unstained WT and fluorescent-minus-one (FMO) controls. Sorted cells (AriaII/Fusion/FlowJov10, BD) were collected in 50% FBS/PBS.

### Hematopoietic assays

YS, AGM, ESCs or LSK-SLAM FL cells were cultured in Methylcellulose (M3434, StemCellTechnologies; 37°C, 5%CO_2,_ 10 days) and CFU-C scored. LSK-SLAM FL cells (1 FL/recipient or 3, 10 or 30 cells/recipient) were injected intravenously into recipients (2×4.5 Gy γ-irradiation). After 4, 16 and 23 weeks, peripheral blood was analysed by CD45.1/CD45.2-flow cytometry. Multilineage analysis on tissues and primary LSK BM cells injected into irradiated secondary recipients were performed at 23 weeks. Recipients with ≥5% donor-derived cells were considered reconstituted.

### Molecular analyses

DNA was extracted from individual CFU-C, ear notches or sorted cells in PCR Buffer (50mM KCl, 10mM TrisHCl, pH8.3, 2.5mM MgCl_2_, 0.1mg/ml gelatin, 0.45%(v/v) Igepal/NP40, 0.45%(v/v) Tween20)+ProteinaseK (10mg/ml, 1h, 55°C) and heat-inactivated (95°C, 10’). 1µl used for *Gpr56* PCR.

RNA was isolated (RNeasyMicroKit, Qiagen), cDNA synthesized (oligodT, Invitrogen; SuperScriptIII, LifeTechnologies) and RT-PCR performed with FastSybrGreen (LifeTechnologies) and primers (**Table EV2**).

Western blotting: 10^6^ ESCs were washed (2xPBS), re-suspended in ice-cold RIPA buffer+protease+phosphatase inhibitors (ThermoFisher), incubated (30’, ice), sonicated, centrifuged (maximum speed, 15’) and protein quantified (BSA kit, BioRad). Samples were boiled (95°C, 5’, SDS buffer, BioRad), SDS-polyacrylamide gel (NuSTep) separated, transferred (30’, 20V) to nitrocellulose (Amersham). Membranes were blocked (5% semi-skimmed milk:TBS-Tween20), blotted (ON; 4°C, 1.5% for Gpr56; RT, 2hr for β−actin; **Table EV1**) and washed (3×5’, TBS-Tween), blotted with HRP-conjugated secondary antibodies (RT, 1hr) and analysed by OdysseyFC (Li-Cor) using ImageStudioLite.

### ESC culture

IB10 ESCs (*WT, G2V, G2V*.*56*^*KO*^, *G2V*.*56*^*KO*^:*97*^*KO*^) were cultured (37°C, 5%CO_2_) on irradiated mouse embryonic fibroblasts (MEFs) in ES medium; DMEM(Lonza), 15% FCS(HyClone), 2mM GlutaMAX, 1mM Na-pyruvate, 1%P/S, 50mM *ß-*mercaptoethanol (all Gibco), 0.1mM non-essential amino acids(Lonza), 1,000U/mL LIF(Sigma). Cells were trypsinized and MEFs depleted by 30’ incubation in EB medium (IMDM, 15%FCS, 1%P/S. EB induction (40rpm; 25⨯10^3^cells/mL) was in EB medium, 2mM GlutaMAX (Gibco), 50mg/mL ascorbic acid(Sigma), 4 ⨯ 10^−4^M monothioglycerol(Sigma), 300mg/mL transferrin(Roche) and supplemented (d3) with 5% proteome-free hybridoma medium(Gibco). From d6, 100ng/mL SCF, 1ng/mL IL-3, 5ng/mL IL-11(Peprotech) and from d4, 10μM Y-27632 ROCK inhibitor and 0.84nM Collagen3a were added. Reagents were refreshed every other day.

### Zebrafish

Zebrafish (*Danio rerio*) embryos were raised at 28.5^°^C(Solaimani Kartalaei et al., 2015). Heterozygous *6*.*0itga2b:EGFP* (CD41-GFP(Lin, Traver et al., 2005)) and *- 0*.*8flt1:RFP* (Flt1:RFP)(Bussmann, Bos et al., 2010) zebrafish were maintained by crosses with WT. Embryos imaged on a LeicaSP5. All mRNA *in vitro* expression constructs were generated by amplification of cDNA with specific primers (**Table EV2**). PCR products were cloned(*p-GEM-T)*, sequence verified and mRNA generated (Ambion sp6/T7 kit). Antisense morpholino against the splice site of the second intron (**Table EV2**; Gene Tools) was dissolved in MQ to1mM (1nl MO was injected in a 1/5 concentration in 0.1M KCl and phenol red). For mRNA rescue experiments, 1nl of 50ng/μl mRNA and 200μM was injected.

### *G2V*.*56*^*KO*^ and *G2V.56* ^*KO*^*/97* ^*KO*^ ESCs

Mouse *G2V.Gpr56* and *Gpr97* knockout ESCs were generated with *CRISPR/Cas9*(Ran, Hsu et al., 2013). Guide(g)RNAs (**Table EV2**) (http://www.e-crisp.org/E-CRISP/designcrispr.html) cloned into *pSp-Cas9(BB)-2A-GFP* (Addgene:48138). Mouse *G2V* ESCs(Kaimakis et al., 2016, Kauts et al., 2018) were transfected (DreamFect, OZBiosciences), GFP^+^ cells seeded on MEFs at 48h, expanded and screened.

### Statistical Analysis

All graphs were generated using GraphPadPrism. One-way ANOVA corrected by Bonferroni’s test was used to compare >2 groups and the Student-t-test for 2-group comparisons. The Fisher exact test determined non-random associations in lineage-bias analysis. *p≤0.05, **p≤0.01, ***p≤0.001.

## Acknowledgements

The authors thank A. Rossi, F. Glykofrydis and lab members for assistance and critical discussions concerning this study. We acknowledge the assistance of the QMRI flow & cell sorting facility (S. Johnston, W. Ramsay, M. Pattison) and the BVS service staff. These studies were supported by ERC AdG (341096), Blood Cancer UK (18010), ZonMW TOP (91211068) and NIH (RO37 DK54077).

## Authorship Contributions

E.D., A.M., S.A.M. and E.P. conceived the project and designed the experiments. E.D. and S.A.M directed project. A.M., S.A.M. and E.P. performed the experiments. X.P. provided the *loxGpr56* mice. C.S.V. and S.A.M. performed hematopoietic transplantation experiments and A.M. and C.S-R. performed in vitro culturing and HPC assays. M.L.L helped generating the *Gpr56*^*KO*^ ES cells. A.M. and S.A.M. generated *CRISPR/Cas9* constructs. A.M., S.A.M. and C.S-R. performed molecular analyses.

E.D., A.M. and S.A.M. wrote the manuscript.

## Expanded View Figure Legends

**Table EV1.**
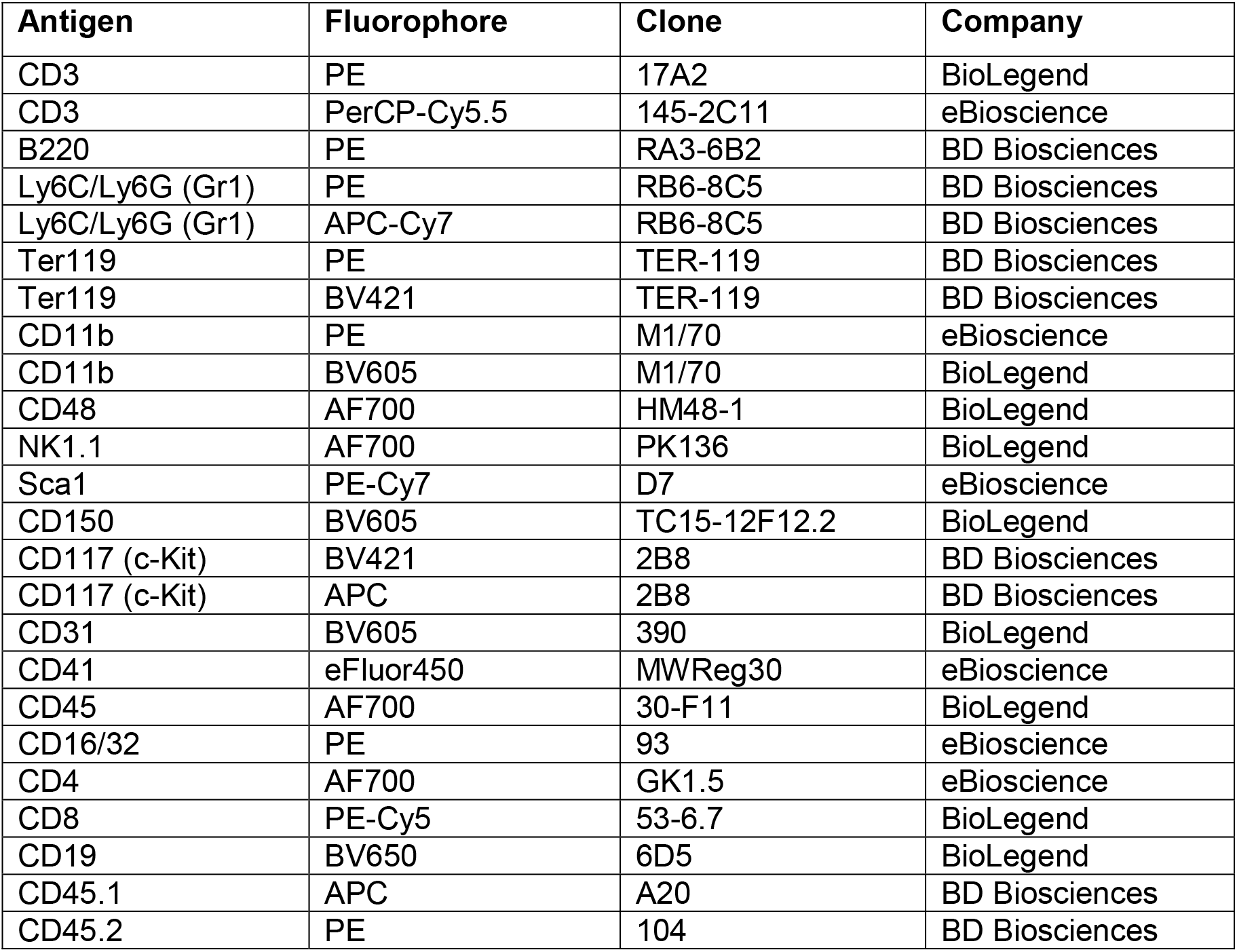
Antibodies.

| **Antigen** | **Fluorophore** | **Clone** | **Company** |
| --- | --- | --- | --- |
| CD3 | PE | 17A2 | BioLegend |
| CD3 | PerCP-Cy5.5 | 145-2C11 | eBioscience |
| B220 | PE | RA3-6B2 | BD Biosciences |
| Ly6C/Ly6G (Gr1) | PE | RB6-8C5 | BD Biosciences |
| Ly6C/Ly6G (Gr1) | APC-Cy7 | RB6-8C5 | BD Biosciences |
| Ter119 | PE | TER-119 | BD Biosciences |
| Ter119 | BV421 | TER-119 | BD Biosciences |
| CD11b | PE | M1/70 | eBioscience |
| CD11b | BV605 | M1/70 | BioLegend |
| CD48 | AF700 | HM48-1 | BioLegend |
| NK1.1 | AF700 | PK136 | BioLegend |
| Sca1 | PE-Cy7 | D7 | eBioscience |
| CD150 | BV605 | TC15-12F12.2 | BioLegend |
| CD117 (c-Kit) | BV421 | 2B8 | BD Biosciences |
| CD117 (c-Kit) | APC | 2B8 | BD Biosciences |
| CD31 | BV605 | 390 | BioLegend |
| CD41 | eFluor450 | MWReg30 | eBioscience |
| CD45 | AF700 | 30-F11 | BioLegend |
| CD16/32 | PE | 93 | eBioscience |
| CD4 | AF700 | GK1.5 | eBioscience |
| CD8 | PE-Cy5 | 53-6.7 | BioLegend |
| CD19 | BV650 | 6D5 | BioLegend |
| CD45.1 | APC | A20 | BD Biosciences |
| CD45.2 | PE | 104 | BD Biosciences |

**Table EV2.**
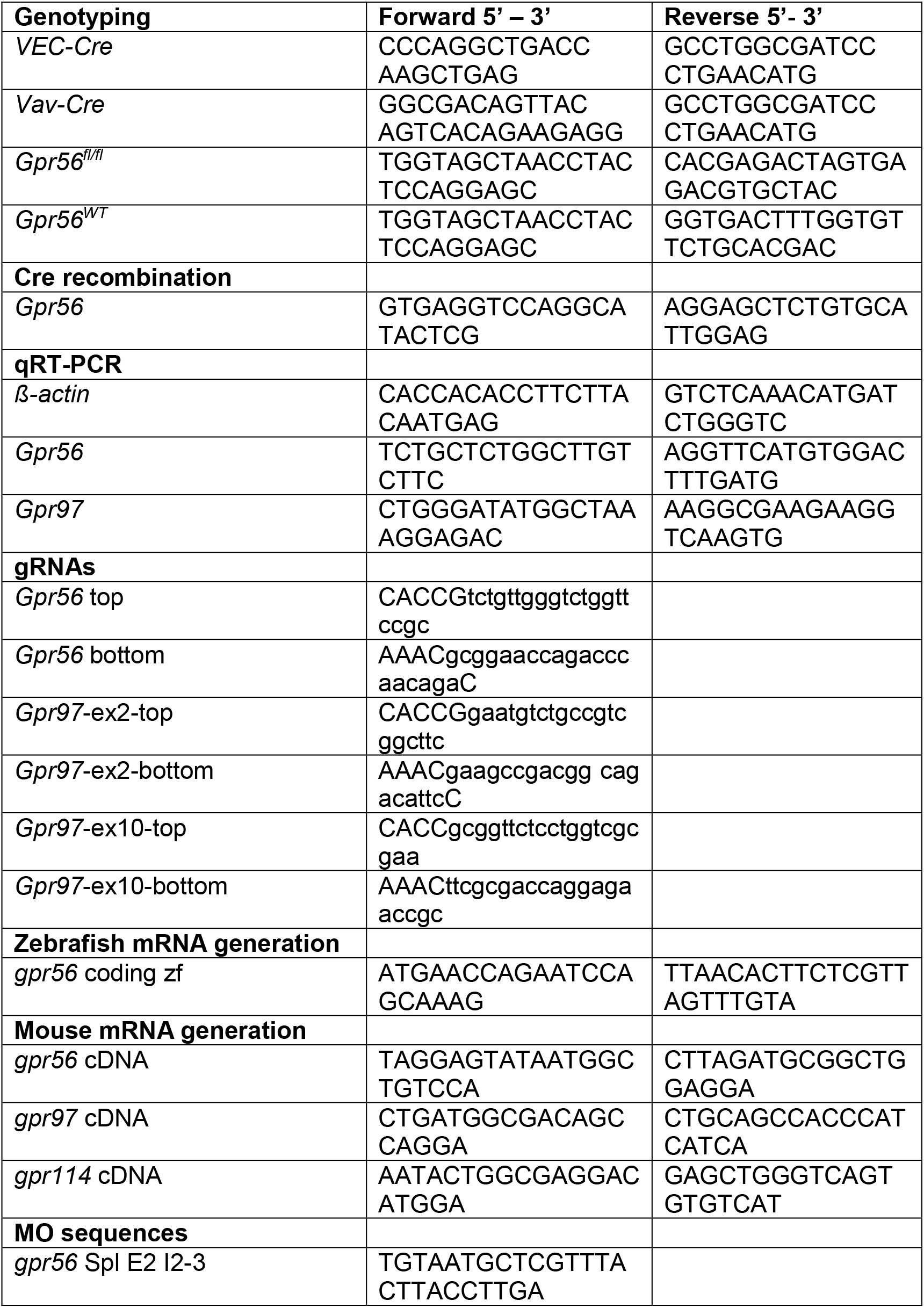
Primers and g-RNAs.

**Figure EV1.**
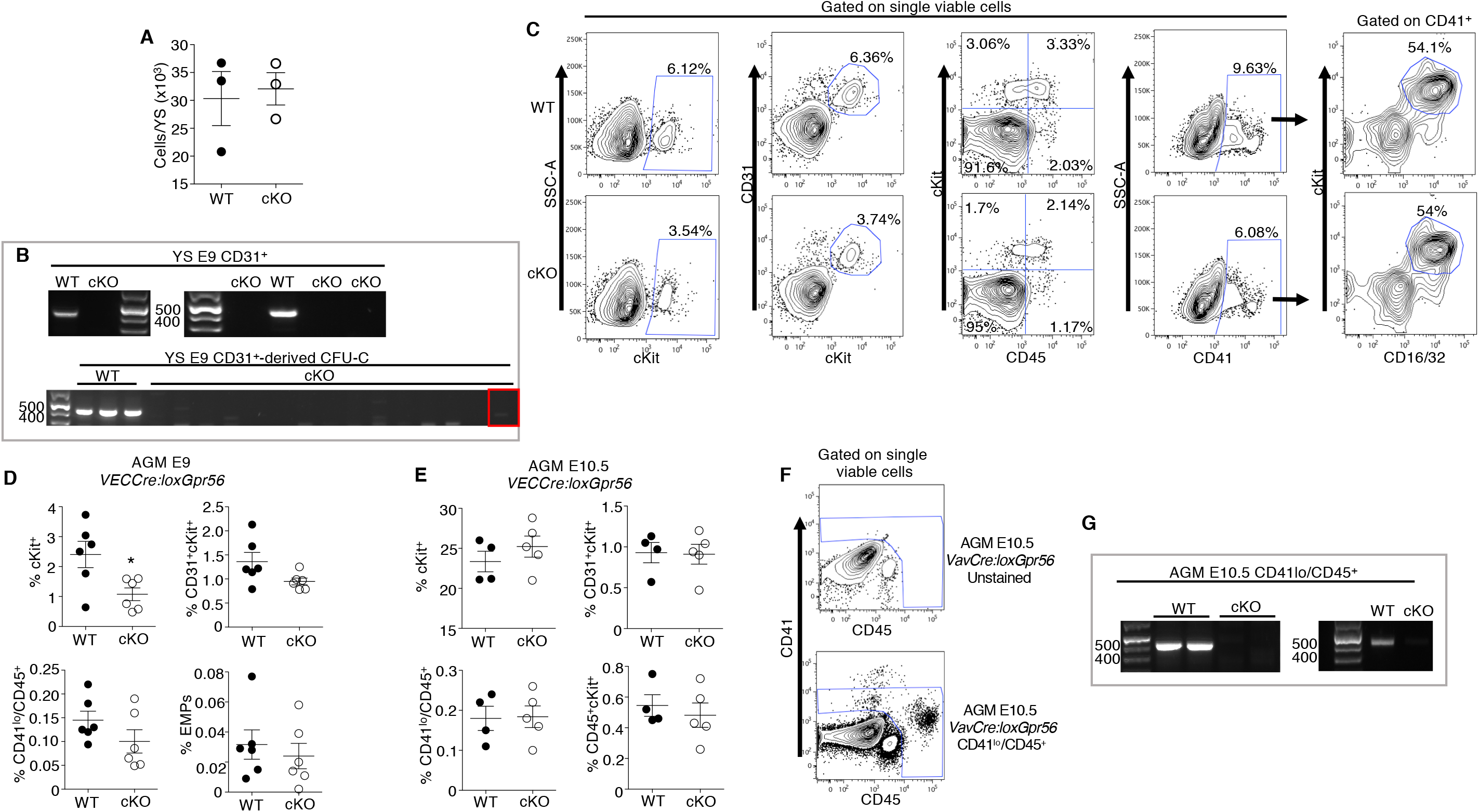
Gating strategy for progenitor analysis and Gpr56 deletion in YS and AGM. A) Number of cells in WT and *VECCre:loxGpr56* cKO E9 YS. WT=black; cKO=white. B) DNA PCR showing the deletion of Gpr56 in sorted CD31^+^ WT control and *VECCre:loxGpr56* cKO E9 YS cells (top panel) and in WT and *VECCre:loxGpr56* E9 YS CD31-derived CFU-C (bottom panel). WT band=460bp. C) Contour plots showing the gating strategy for the progenitor analyses in Fig 1D and Fig1H. One WT and one E9 YS *VECCre:loxGpr56* are shown as representative examples. D) Percentages of cKit^+^, CD31^+^cKit^+^, CD41lo/CD45^+^ and CD41^+^cKit^+^CD16/32^+^ (EMP=erythromyeloid progenitor) cells in WT (black) and cKO (white) E9 AGM (n=6). E) Percentages of cKit^+^, CD31^+^cKit^+^,CD41^lo^/CD45^+^ and CD45^+^cKit^+^ cells in WT (black) and cKO (white) E9 AGM (WT n=4, cKO n=5). F) Contour plots showing the gating strategy for the cell sorting of CD41^lo^/CD45^+^ from E10.5 *VavCre:loxGpr56* AGM. G) DNA PCR showing the deletion of Gpr56 in sorted CD41lo/CD45+ WT control and *VavCre:loxGpr56* cKO cells from E10.5 AGM. WT band=460bp.

**Figure EV2.**
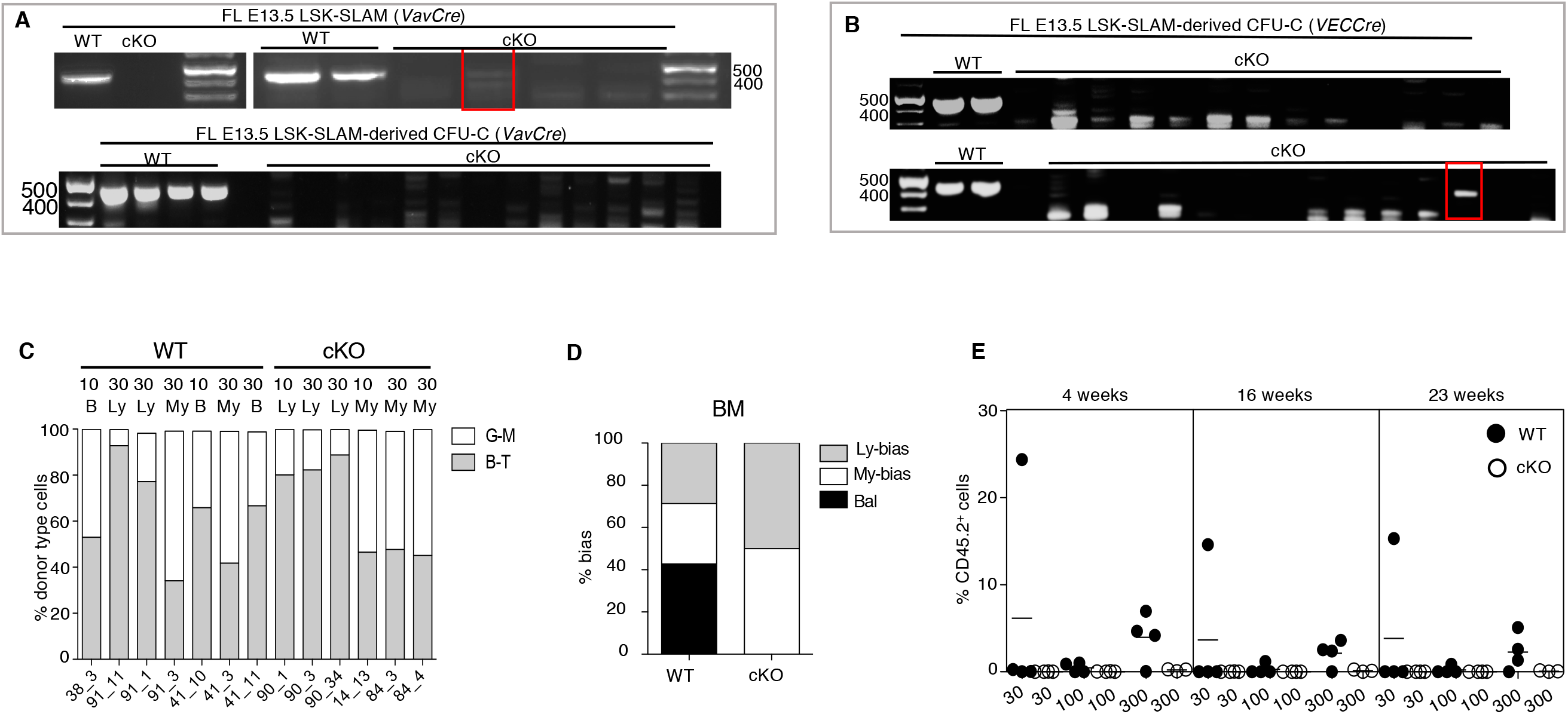
Gpr56 is deleted upon Cre activation in E13.5 FL influencing HSC lineage bias and the self-renewal. A) DNA PCR showing the deletion of *Gpr56* in sorted LSK-SLAM cells from WT and *VavCre:loxGpr56* cKO E13.5 FL (top panel) and in LSK SLAM *VavCre:loxGpr56* FL-derived CFU-C (bottom panel). WT band=460bp. B) DNA PCR showing the deletion of *Gpr56* in sorted LSK-SLAM cells from E13.5 *VECCre:loxGpr56* FL-derived CFU-C. WT band=460bp. C) Percentage of lymphoid and myeloid cell contribution in BM of 13 individual adult irradiated recipient mice as measured by flow cytometry at 23 weeks post-injection of 10 and 30 LSK-SLAM WT control and *VavCre:loxGpr56* cKO E13.5 FL cells. D) Percentages of lymphoid-biased, balanced and myeloid-biased HSC engrafted recipients from panel C. E) Donor cell-derived chimerism of PB of secondary recipient mice at 4, 16 and 23 weeks post-transplantation of bone marrow LSK cells from primary recipients of WT and *VavCre:loxGpr56* cKO sorted E13.5 FL cells. Percentage of CD45.2+ donor cells in PB is shown. Horizontal lines indicate average engraftment percentage. WT=black; cKO=white. n=4 per group.

**Figure EV3.**
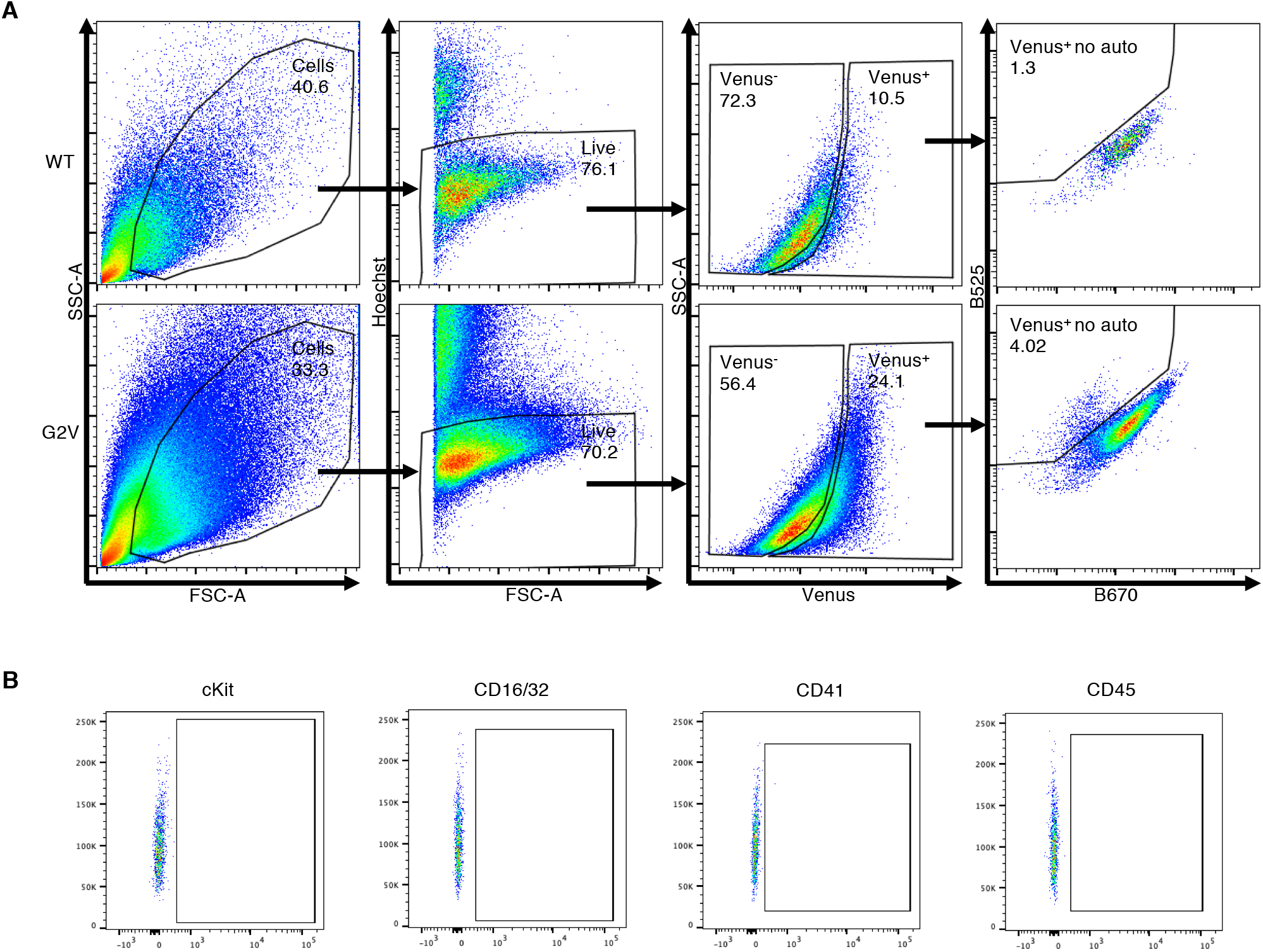
Gating strategy for WT and G2V ESC differentiation cultures. Forward scatter (FSC), side scatter (SSC), Venus, and Venus minus autofluorescence plots are shown in upper panels. After gating on live cells, the Venus^+^ cells are plotted against a dump neighbour channel (B525-Venus vs B670) to exclude auto-fluorescent cells. Only the cells positive in the B525 channel and not in the B670 channel are considered Venus^+^. WT cells are used as control to set the gates. The Fluorescent minus one (FMO) controls for cKit, CD16/32, CD41, CD45 antibodies are shown in lower panels.

**Figure EV4.**
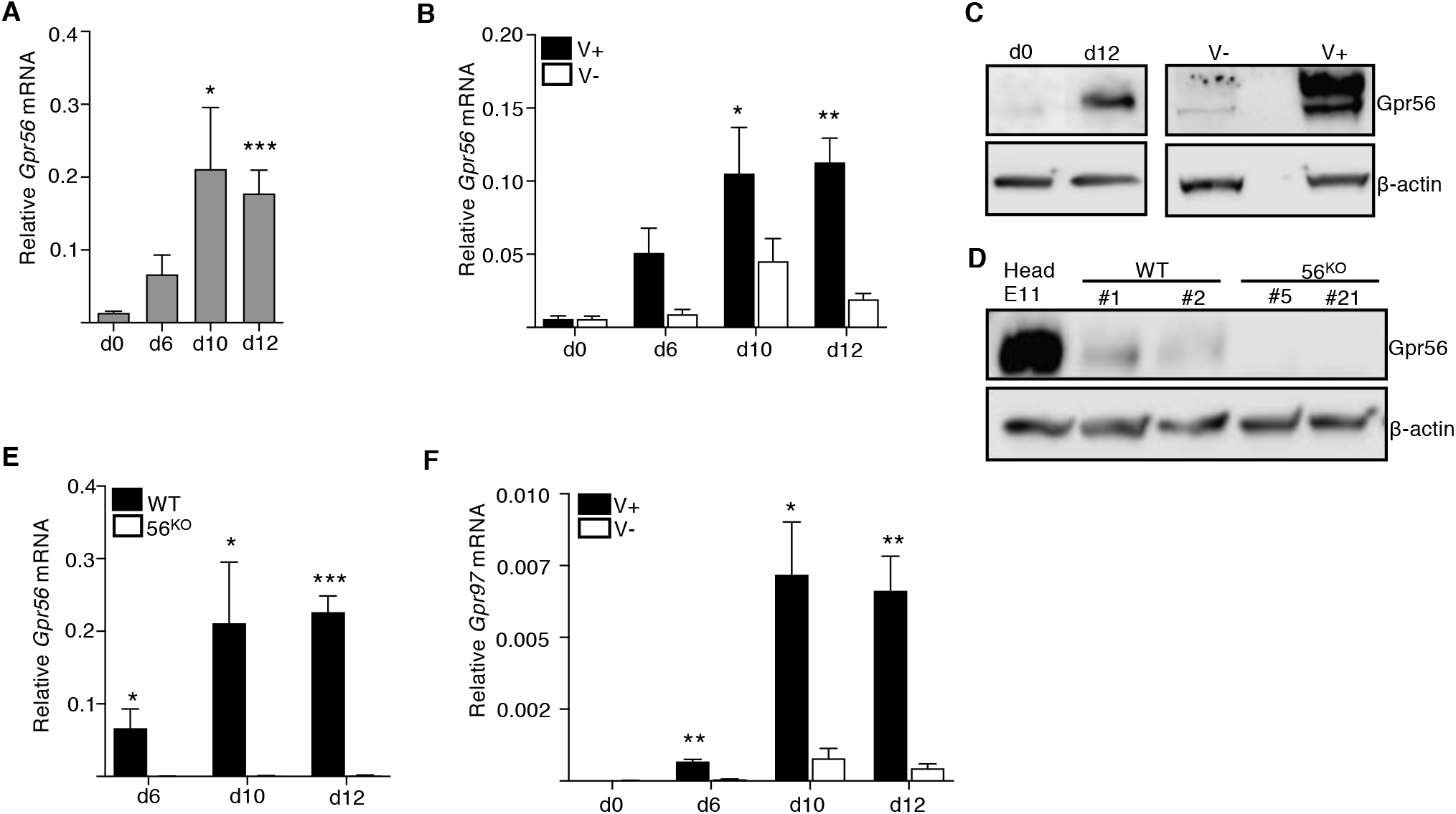
Gpr56 and Gpr97 are expressed in mouse G2V.WT differentiated cells. Time course qRT-PCR analysis of relative *Gpr56* expression (normalized to *β-actin*) A) in unsorted and B) in Venus sorted G2V ESC hematopoietic differentiation cultures. C) Western blot analysis of Gpr56 protein expression in day 0 and day 12 unsorted G2V ESC hematopoietic differentiation cultures (left panel) and day 12 Venus sorted G2V ESC hematopoietic differentiation cultures (right panel), with β-actin as protein normalization control. D) Western blot analysis of Gpr56 protein expression in day 6 G2V.WT (clones #1 and #2) and G2V. 56KO (clones #5 and #21) ESCs, with β-actin as protein normalization control. Mouse embryonic head used as positive control. E) Time course qRT-PCR analysis of relative *Gpr56* expression (normalized to *β-actin*) in unsorted G2V.WT and G2V. 56KO (clone #5) ESC hematopoietic differentiation cultures. F) Time course RT-qPCR analysis of relative *Gpr97* expression (normalized to β-actin) in G2V.WT Venus sorted cells. d=day of culture harvest; *p<0.05: **p<0.01; ***p<0.001.

**Figure EV5.**
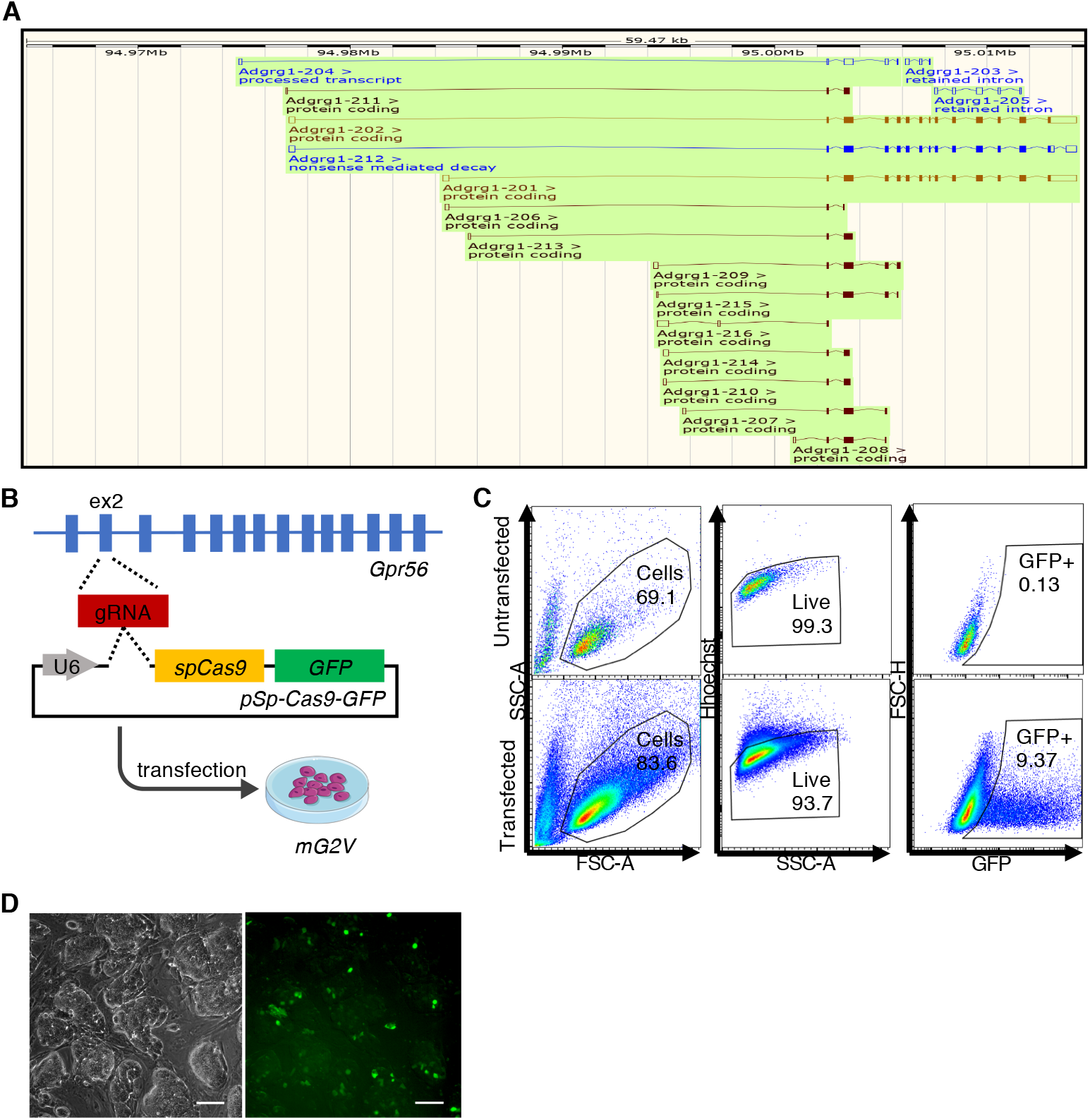
Generation of *Gpr56* deleted G2V ESCs. A) *Gpr56* gene and splice isoforms as taken from Ensemble database. B) CRISPR/Cas9 strategy showing gRNA for *Gpr56* exon 2 and insertion in pSp-Cas9-2A-GFP vector used for transfection of G2V ESCs. C) Flow cytometric analysis of transfected and untransfected control ESCs for forward (FSC) and side scatter (SSC), viability and GFP expression. D) Fluorescence microscopic images of untransfected and transfected ESCs at 48 hours post-transfection.

**Figure EV6.**
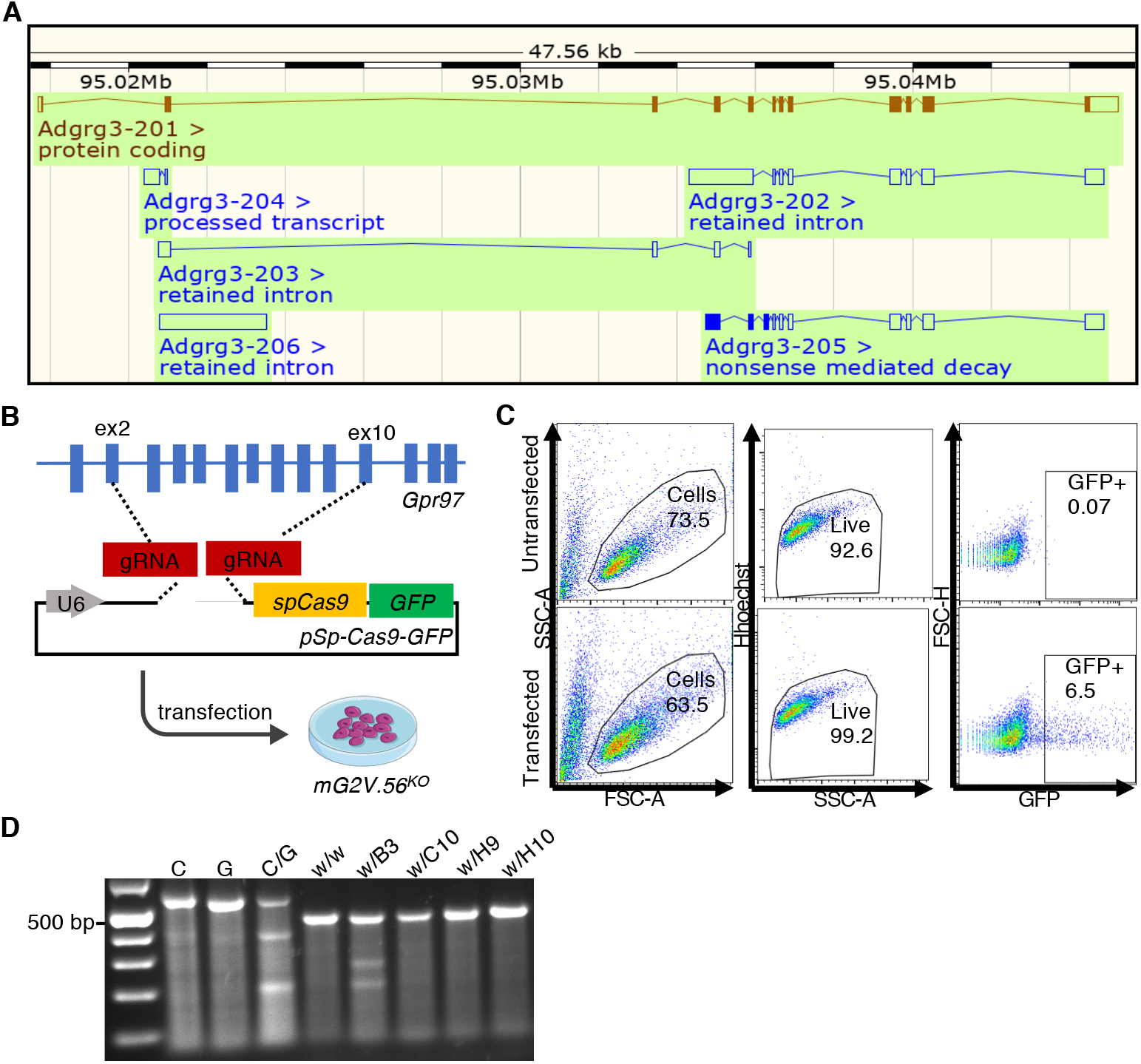
Generation of *Gpr56:Gpr97* double deleted G2V ESCs. A) Mouse *Gpr97* gene and splice isoforms as taken from Ensemble database. B) CRISPR/Cas9 strategy showing gRNAs for *Gpr97* exons 2 and 10 and insertion in pSp-Cas9-2A-GFP vector used for transfection of G2V.56KO ESCs. Two rounds of transfection were performed, first with exon 2 gRNA, and after sorting for GFP^+^ cells a second transfection was performed with exon 10 gRNAs. C) Flow cytometric analysis of transfected and untransfected control ESCs for forward and side scatter, viability and GFP expression at 48 hours post-second transfection. D) Surveyor assay on DNA from 4 CRISPR/Cas9 ESC clones (B3, C10, H9, H10; negative for *Gpr97* mRNA) after second transfection and sorting. Clone B3 shows a mismatch in *Gpr97* genomic sequence. C and G are negative and C/G is positive controls.

